# Analysis of continuous infusion functional PET (fPET) in the human brain

**DOI:** 10.1101/778357

**Authors:** Shenpeng Li, Sharna D. Jamadar, Phillip G.D. Ward, Malin Premaratne, Gary F. Egan, Zhaolin Chen

## Abstract

Functional positron emission tomography (fPET) is a neuroimaging method involving continuous infusion of 18-F-fluorodeoxyglucose (FDG) radiotracer during the course of the PET examination. Compared with the conventional bolus administered static FDG PET which provides only a snapshot of the averaged glucose uptake into the brain in a limited dynamic time window, fPET offers a significantly wider time window to study the dynamics of glucose uptake. Several earlier studies have applied fPET to investigate brain FDG uptake and study its relationship with functional magnetic resonance imaging (fMRI). However, due to the unique characteristics of fPET signals, modelling of the fPET signal is a complex task and poses challenges for accurate interpretation of the results. This study applies independent component analysis (ICA) to analyze resting state fPET data, and to compare the performance of ICA and general linear modelling (GLM) for estimation of brain activation in response to tasks. The fPET signal characteristics were compared using GLM and ICA methods to model the fPET visual activation data. Our aim was to evaluate GLM and ICA methods for analyzing task fPET datasets and present ICA method in the analysis of resting state fPET datasets. Using both simulation and *in-vivo* experimental datasets, we show that both methods can successfully identify task related brain activation. We report fPET metabolic resting state brain networks analyzed using the fPET ICA method in a cohort of healthy subjects. Functional PET provides a unique method to map dynamic changes of glucose uptake in the resting human brain and in response to extrinsic stimulation.

## 1. Introduction

Positron emission tomography (PET) is a powerful neuroimaging method to measure molecular processes *in vivo* in the living human brain. 18-F-fluorodeoxyglucose (FDG) is a widely used PET radiotracer for quantitatively measuring cerebral glucose uptake, which can be used to infer glucose metabolism in brain imaging studies (Phelps et al. 1979; Reivich et al. 1985). As one of major energy sources of the brain, the dynamics of glucose metabolism are highly correlated with the fluctuations of brain activity (Figley and Stroman 2011). The static FDG PET with conventional bolus injection measures the average glucose utilization throughout the scan for task (Kushner et al. 1988) or resting state studies (Savio et al. 2017). Recent efforts have been made to explore the dynamics of brain glucose metabolism by multi-frame PET reconstruction following a bolus injection (Jamadar, Ward, Carey, et al. 2019; Passow et al. 2015; Tomasi et al. 2017). However, the information of dynamic change of FDG uptake using bolus administration is limited due to lack of continuous supply of FDG. During 1990s, the idea of continuous infusion was pioneered by Carson and others to demonstrate accurate receptor measurements (Carson 2000; Carson et al. 1993). Recently, Villien et al. (2014) applied continuous infusion of [18-F]FDG to identify dynamic glucose uptake in the brain, and named the technique ‘functional PET’ (fPET). The aim of the fPET approach is to maintain a continuous plasma supply of FDG to provide improved sensitivity of brain-state changes, and better temporal dynamics than bolus PET to track dynamic change of glucose uptake over time. Several studies have demonstrated that fPET can isolate task related changes in glucose uptake (Hahn et al. 2016; Hahn et al. 2018; Jamadar, Ward, Li, et al. 2019; Rischka et al. 2018; Villien et al. 2014; Jamadar, Ward, Carey, et al. 2019).

The development of data analysis methods is critically important for correct interpretation and inference of neuroimaging findings. There are two major approaches to analyze functional brain data: model-based approaches and data-driven methods. The general linear model (GLM) is a the most common model-based method using in neuroimaging that interprets each voxel in 4D functional images as a linear combination of task design regressors and nuisance variables. The estimated parameters are then submitted to parametric statistical tests to generate brain activation maps (Friston 2007). The GLM model-based approach has been widely used in the analysis of functional MRI (fMRI), EEG and recently in fPET (Hahn et al. 2016; Villien et al. 2014; Jamadar, Ward, Carey, et al. 2019). In the fPET GLM analysis, the time activity signal is represented by the combination of the task regressors, baseline regressor, and other nuisance regressors accounting for motion and physiological noises (Hahn et al. 2016; Villien et al. 2014). Task-related activated brain regions can be identified using the GLM method.

Compared with model-based methods, data-driven analysis methods such as independent component analysis (ICA) explore the intrinsic coherence of the data rather than fitting an explicit model. ICA has been widely applied to resting state fMRI (Beckmann et al. 2009; Beckmann et al. 2005; Beckmann and Smith 2004; Calhoun et al. 2001a; Damoiseaux et al. 2006) and task-based fMRI (Calhoun et al. 2006; Calhoun et al. 2001b; McKeown, Hansen, and Sejnowsk 2003; McKeown et al. 1998; McKeown and Sejnowski 1998). We have previously applied ICA in a visual task fPET experiment to investigate its relationship with simultaneously acquired fMRI data (Jamadar, Ward, Li, et al. 2019). In resting state PET, ICA has also been applied to bolus-administered FDG PET to investigate brain glucose metabolism and connectivity (Savio et al. 2017; Tomasi et al. 2017).

Modelling of the fPET signal is similar to modelling other functional neuroimaging data such as fMRI and electroencephalogram (EEG). However, compared with fMRI in particular, fPET signals have several distinct characteristics. Firstly, fMRI measures approximately the blood-oxygen-level-dependent (BOLD) signal at each instantaneous temporal sampling point, and the task related BOLD response periodically fluctuates around a baseline. By contrast, fPET signals monotonically increase over time and measure the accumulated FDG from the beginning of the administration of the tracer to the current acquisition time. Therefore, fPET measures a response integrated over time. Secondly, the temporal resolution of fPET signals is lower than fMRI (e.g. 1 min vs 2 secs), because of the fundamental limitation of the FDG PET count statistics and the PET detector technology (Chen et al. 2018). Thirdly, the BOLD response has been extensively studied, e.g., the canonical haemodynamic response function (HRF) (Friston et al. 1998; Friston, Jezzzard, and Turner 1994; Lindquist et al. 2009; Rajapakse et al. 1998) and balloon models (Buxton, Wong, and Frank 1998; Friston et al. 2000), whereas the neural basis of real time glucose uptake has not been fully understood due in part to the limited availability of experimental data.

In this paper we have demonstrated the application of the data-driven ICA method for analyzing resting state fPET datasets, and compared the performance of GLM and ICA for estimation of task fPET brain activations. The aim of the study was to evaluate GLM and ICA methods for analyzing task fPET datasets and present ICA method in the analysis of resting state fPET datasets. We investigated fPET data modelling and interference procedures in both GLM and ICA approaches. Using a simulated task fPET experiment, GLM and ICA methods were quantitatively compared and validated. GLM and ICA methods were then applied to an *in-vivo* visual task fPET dataset for comparison and interpretation. Finally, we apply the optimized ICA method to analyze resting state fPET data in a cohort of 28 of healthy subjects. We hypothesised that resting brain metabolic networks can be obtained using the data driven ICA approach.

## 2. Materials and Methods

The section provides an overview of methods for analysis of fPET activity signals and brain activation mapping, and an overview of the model-based GLM method (Figure 1a) and the data-driven ICA method (Figure 1b).

**Figure 1:**
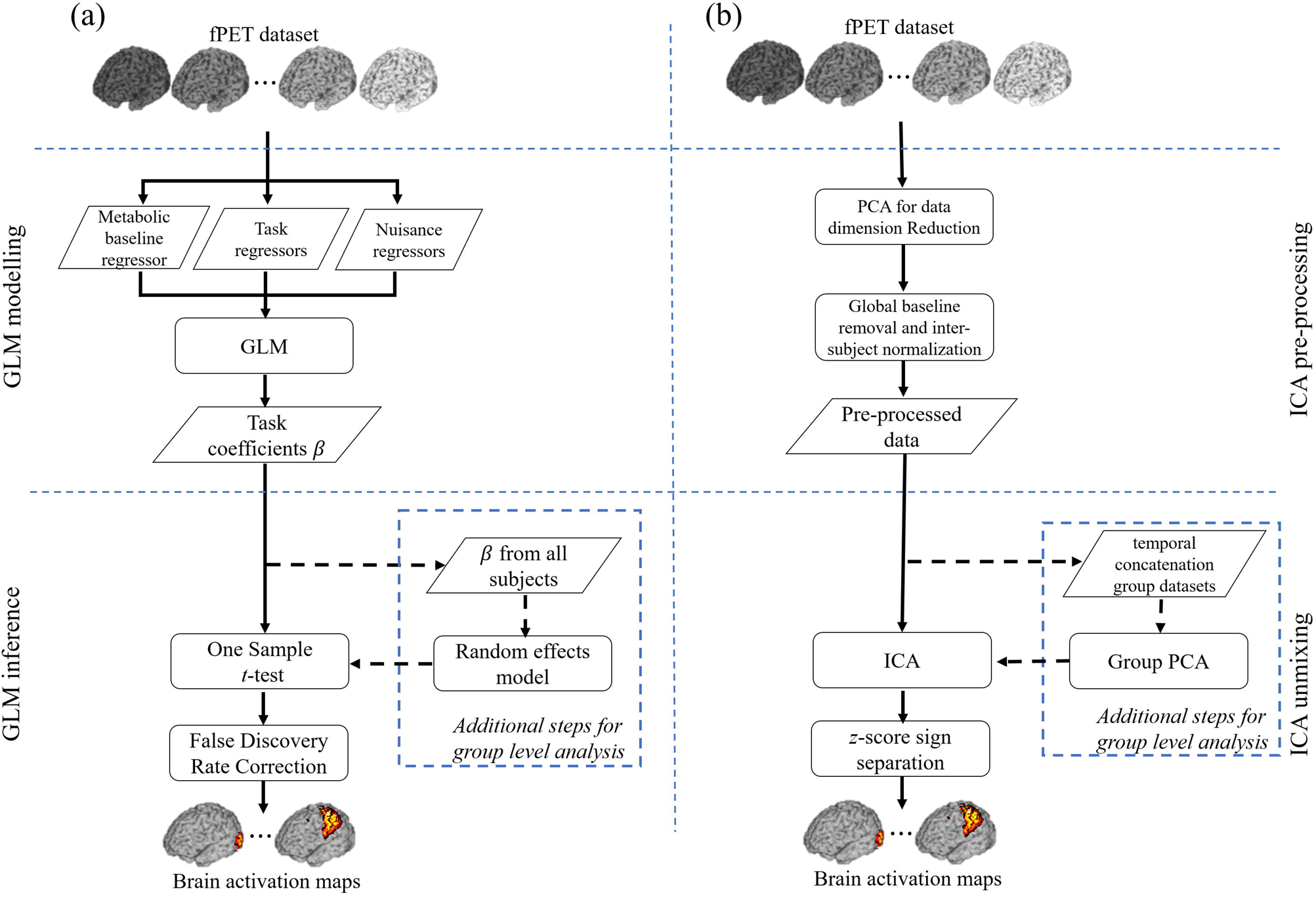
Overview of fPET data analysis methods (rectangles represent a process and parallelograms represent input/output datasets), **(a)** GLM, and **(b)** ICA.

### 2.1 GLM of fPET datasets

Using GLM, the FDG time activity signal can be modelled as a linear combination of a baseline metabolic signal and a task induced activation signal. The baseline represents a glucose measure that is independent of any task stimulation, and the activation signal is defined as the task induced FDG activity changes (Hahn et al. 2016; Villien et al. 2014). The 4D fPET signal can then be modelled as

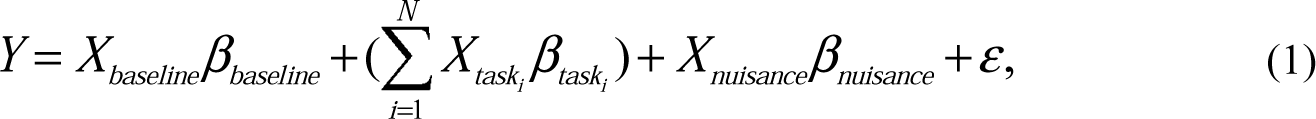

where *Y* is the measured FDG time activity signal at each voxel. *X*s are the design matrices, including a baseline regressor, *X*_*baseline*_, task regressors ranging from *X*_*task*1_ to *X*_*taskN*_ for each task stimulus and a nuisance regressor, *X*_*nuisance*_, which includes head motion, physiological noise and other noise terms. These design matrices incorporate *a priori* knowledge to parameterize the linear model. *β*s are the coefficients to be estimated. The noise term, *ε*, is assumed to be uncorrelated to the measurements.

#### Models of the baseline metabolic regressor and task regressors

Hahn et al. (2016) and Villien et al. (2014) have demonstrated in their work that the fPET signal increases monotonically over time, which is driven by the continuous infusion of FDG into the blood stream at constant rate (36ml/hr). The aim of the baseline regressor is to fit the baseline metabolic signal. In general, the baseline metabolic is a nonlinear function and can be approximated using a polynomial function, e.g. a third-order polynomial (Hahn et al. 2016) that is brain region dependent (Hahn et al. 2016). The task regressors are parameterized as a ramp function with slope being one during stimulation and zero during the resting period, implying an increased FDG uptake rate due to the task stimulation (Hahn et al. 2016; Villien et al. 2014).

As an alternative approach, Villien et al. (2014) noted that the first order derivative of the temporal fPET signals can also be used to fit a GLM. This is because fPET signals are accumulated over time. Using this approach, an *on-off* task design matrix can be applied as in the fMRI GLM analysis. However, in practice, fPET data have poor signal to noise characteristics (i.e. low FDG counts) due to short temporal binning window, such that the derivative operation may dramatically amplify noise.

#### Implementation of GLM analysis for subject level and group level fPET

In single-subject level GLM, multivariate regression was used to estimate the *β* coefficients. One-sample *t*-test against each task coefficient *β*_*task*_ was computed for all voxels to generate the corresponding activation map and the significant level was false discovery rate (FDR) adjusted at the significance level *p* < 0.1 (Benjamini and Hochberg 1995). The activation map was then thresholded at *p* < 0.1. The mean task FDG activity in the activated region is given as *Y* − (*X*_*baseline*_*β*_*baseline*_ + *X*_*nuisance*_*β*_*nuisance*_).

The implementation of group level GLM analysis was based on a two-level analysis approach. In the first level analysis, subjects were first co-registered, and GLM fitting was conducted on each subject to estimate *β* coefficients. Random-effects analysis (Holmes and Friston 1998) was then conducted at the group level. The *β*s weighted by the design contrast coefficients from all subjects were then grouped together to conduct statistical tests (e.g. *t*-test) to derive activation maps during the group inference step (Frison and Pocock 1992; Holmes and Friston 1998).

### 2.2 Spatial ICA of fPET datasets

In contrast to GLM, independent component analysis (ICA) is a full data driven method which has been used widely for both task and resting state fMRI analysis (Calhoun et al. 2001a; McKeown et al. 1998). In the context of fPET, the spatio-temporal data matrix can be represented as a mixture of independent components

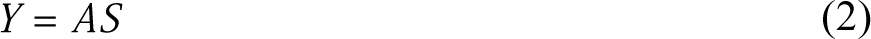

where *Y* is the fPET spatio-temporal data matrix, in which each row contains voxels from each dynamic volume and each column contains FDG time activity signals of the corresponding voxel; *S* is the matrix of independent components, in which each row represents a spatial component, and *A* is the mixing matrix for the corresponding independent components. Both *S* and *A* are unknown. The ICA algorithm aims to estimate *S* by minimizing mutual information or maximizing of non-Gaussianity (Hyvarinen and Oja). The following un-mixing procedure is given by

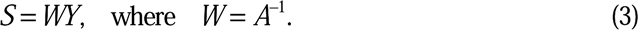

ICA circumvents the requirement for a task design matrix, which is critical in GLM, but intrinsically unknown in resting state experiments. However, ICA requires pre-processing of fPET signals to remove the global baseline signal (i.e. whole brain averaged FDG update) for the accurate estimation of spatial independent components.

#### Removal of global baseline fPET signal

Unlike GLM, which requires of an accurate model fit of the baseline metabolic signal, ICA relies on the statistical distribution of signals themselves (Hyvarinen and Oja 2000). However, the baseline FDG uptake in the brain accounts for a large proportion of the overall signal variance in the fPET signals, and this should be removed before the ICA unmixing step to improve the sensitivity for calculating the spatially independent components. The global baseline can be removed by spatially normalizing the data to remove the mean in each voxel and dividing the voxel wise standard deviation of the volume. This is equivalent to *z*-scoring the signals, which converts the fPET signal into a regionally relative signal. Following this normalization, the brain networks resulting from the ICA method can be interpreted in the context of coherent relative signal changes. Importantly, care should be taken, particularly in task-based analysis, where the global subtraction of the mean signal can introduce seemingly spurious negative correlations.

#### Implementation of single-subject spatial ICA of fPET data

To implement the spatial ICA method on a single subject, a grey matter mask including both cerebral and cerebellar grey matter was applied on the 4D fPET data and the masked signals were then reshaped into a *t* × *n* (*t* < *n*) spatio-temporal matrix *Y*, where the row of *Y* represent the vectorisation of masked volume at each timepoint, and the column of *Y* the temporal sequence of each voxel. Principal component analysis (PCA) was applied on *Y* to generate the dimension-reduced data, *Y*_*D*_ *= E*^*T*^*Y*, where *E* is orthonormal reduction matrix with size of *t*_*D*_ × *t, t*_*D*_ *< t* (Jolliffe and Cadima 2016). Then, *Y*_*D*_ was projected back to the original dimension 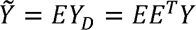. The rank of 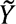 is *t*_*D*_ and it approximates the original signal. For every row of 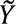, the global baseline was removed, as described previously. The normalized data 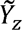 was then decomposed by an ICA unmixing algorithm (e.g. FastICA (Hyvarinen and Oja 1997) or InfoMax (Bell and Sejnowski 1995)) to extract the spatial components *S*. They were then converted to the standard score (*z*-score), and each component was then separated into a positive segment, *S*_*P*_, and a negative segment, *S*_*N*_, to calculate their corresponding timecourse. For each positive and negative activation, the corresponding timecourse was computed from the normalized data 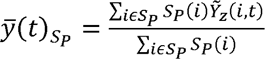, and 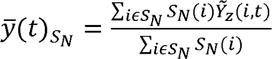, respectively.

#### Implementation of group-level spatial ICA of fPET data

Group-level spatial ICA can be performed using the temporal concatenation approach, which is widely used in group fMRI analysis (Calhoun et al. 2001a). All subjects were co-registered using the MNI-152 template. A grey matter mask from the MNI structural atlas (Mazziotta et al. 2001) and probabilistic cerebellar atlas (Diedrichsen et al. 2009) were used to mask and reshape the 4D fPET data to spatio-temporal matrix for each subject.

One important factor for group fPET analysis is to remove inter-subject data variability which is caused by FDG dose variability, subject physiology, and other experimental factors. The procedure of global baseline removal also removed inter-subject data variability. Theoretically, ICA is robust to inter-subject variability including signal amplitude, as ICA fundamentally unmixes a signal based on the probability distribution of the signal. A pre-whitening step prior to the ICA decomposition can remove such a variation and speed up the convergence for ICA (Hyvarinen and Oja 2000). However, in practice, we employ an additional group level PCA to reduce data dimensionality before conducting the group level ICA. The group level PCA is based on the variance of the signal cross subjects, with larger weighting for data with greater variance. Therefore, the *z*-scoring step from the global baseline removal is performed for each subject to normalize inter-subject variability.

The global baseline removed data matrix from all subjects were temporally concatenated to from a group spatio-temporal matrix. A group-level PCA was applied to further reduce computational complexity. ICA was used to extract group level *z*-scored spatial components. The same procedure described in the single-subject ICA can be followed to calculate timecourses for each subject by using the group component. Finally, the group level timecourse was derived as the mean of timecourses from all subjects.

Once group level independent component maps were calculated, subject-specific independent component maps were derived by a dual regression approach (Beckmann et al. 2009). To achieve this, a subject-specific timecourse was estimated by regressing the group level component with the normalized fPET dataset for each subject. Then, the second regression was performed by using obtained timecourse to generate subject-specific component maps.

### 2.3 Simulation Experiments

To validate both GLM and ICA methods for application in fPET experiments, we designed a synthetic experiment which included the simulation of kinetic model-based brain activation and PET image reconstruction. The kinetic process of FDG was simulated to derive task-based brain activation. The simulated brain activation was regarded as the ground truth, and ICA and GLM were quantitatively compared and evaluated using the simulated data.

#### Brain activation simulation

The FDG metabolism process in the human brain can be modelled as a two tissue compartment model (Phelps et al. 1979; Sokoloff et al. 1977) using the following differential equations

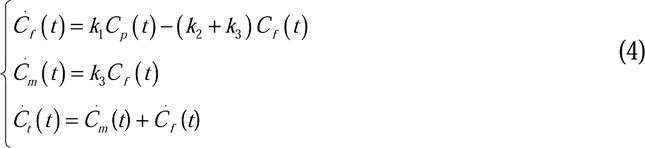

where *C*_*p*_(*t*) is the concentration of FDG in plasma, *C*_*f*_ (*t*) is tissue/unmetabolized FDG concentration, *C*_*m*_(*t*) is the metabolized FDG concentration and *C*_*t*_(*t*) is the total FDG concentration, which is the sum of *C*_*m*_(*t*) and *C*_*f*_(*t*).

The total FDG concentration, *C*_*i*_(*t*), is corrected by blood occupation fraction, *V*_*B*_, which is given by

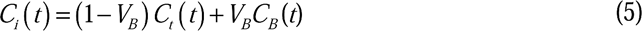

where *V*_*B*_ *=* 0.96 in grey matter and *V*_*B*_ *=* 0.98 in white matter, respectively (Phelps et al.). The concentration in whole blood is given by *C*_*B*_(*t*) = (1 − 0.85*H*)*C*_*P*_(*t*), where *H* = 0.3 is the measured haematocrit for the human brain, and 0.85 is the correction factor (Phelps, Grubb, and Ter-Pogossian 1973). The *C*_*i*_(*t*) in equation (5) is the time activity of the fPET signal based on the given plasma concentration level *C*_*p*_(*t*). The dynamic time activity differences between grey and white matters can be derived using the compartment model in equations (4)-(5) with tissue specific constant rate *k*_l_, *k*_2_ and *k*_3_. We used the constant rate parameters from Table I (Lucignani et al. 1993).

**Table I:** Kinetic constant rates for grey matter and white matter in FDG PET

|  | Grey Matter | White Matter |
| --- | --- | --- |
| $k_1$ (mL/min) | 0.101 | 0.047 |
| $k_2$ (min<sup>-1</sup>) | 0.071 | 0.070 |
| $k_3(\text{min}^{-1})$ | 0.042 | 0.035 |

By substituting the *k*_l_, *k*_2_ and *k*_3_ into equations (4)-(5), the output *C*_*i*_(*t*) was used as the metabolic baseline signal at rest. A 20% increment of *k*_3_ (i.e. a boxcar function) during the task periods was simulated in the activated cortical regions (Villien et al. 2014). Two tasks were considered. The first one was the activation in the visual cortex and the second one was the activation in the left motor cortex. During a 90-min fPET experiment, two sessions of visual stimulation were simulated in the visual cortex. The first one was from 20 mins to 30 mins and the second one started from 70 mins to 80 mins. A single 20-min motor stimulation was simulated during the 40 mins to 60 mins of the fPET experiment.

#### Ground truth GLM regressor

The difference between the kinetic model time activity *C*_*i*_(*t*) with and without change of *k*_3_ is the time activity signal induced by the task. The task induced time activity curve was used as the ground truth GLM regressor in the quantitative comparison between GLM and ICA.

#### PET data simulation and image reconstruction

A T1 weighted Magnetic Resonance (MR) image was used as the template for the simulation experiment. The grey matter, white matter and brain cortex labels were segmented by using Freesurfer with Desikan-Killiany Atlas (Diedrichsen et al. 2009). The segmented regions were masked and co-registered from T1 space to the PET space using the Advanced Normalization Tools (ANTs) (Avants et al. 2011). Figure 2 depicts the brain activation simulation process. The grey matter and white matter time activity signals were simulated based on the kinetic model. The overall grey matter time activity signal was used as the metabolic baseline FDG uptake. The task induced time activity signals in the two task brain regions were simulated from the change of *k*_3_ in the kinetic model. The response of *k*_3_ is not a boxcar function. Both TACs demonstrate the overshoot after the end of task stimulation.

**Figure 2:**
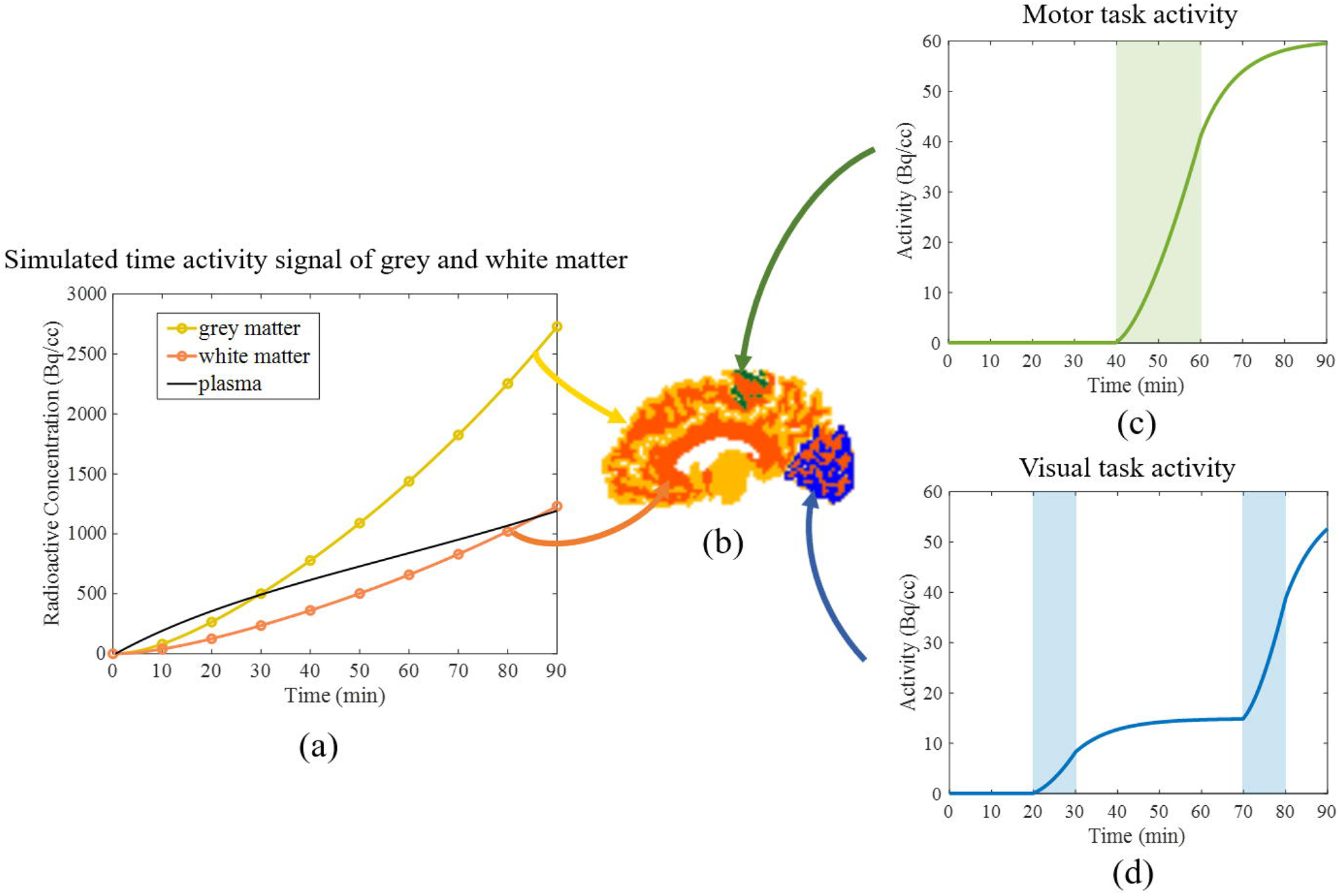
fPET simulation procedures. **(a)**: time activity signals of simulated FDG uptake in grey matter (Yellow), white matter (Red) and plasma input function (Black). **(b):** simulated brain FDG uptake with grey matter mask (Yellow), whiter matter mask (red), motor cortex mask (Green) and visual cortex mask (Blue). **(c)** and **(d):** task induced time activity signals in the motor cortex and the visual cortex. The shadowed areas represent task stimulation periods. The k_3_ parameter has increased by 20% during each task.

For the simulated 90-min dynamic fPET, the duration of each volume was set to one minute, and a total of 90 frames. Each volume was 100 × 100 × 100 voxels with 2 × 2 × 2 mm^3^ resolution. The synthetic PET data acquisition was simulated using the following steps. First, the dynamic image volumes were filtered by a 4 mm FWHM spatial Gaussian filter to simulate the partial volume effect in FDG PET. Then, it was converted to a 344 × 344 × 252 sinogram volume by radon transform with 252 angles in [0, *π*] to simulate the projection of lines of responses (LORs) in PET data acquisition. The thinning Poisson process (Vardi, Shepp, and Kaufman 1985) was used to simulate the low count statistics for dynamic PET reconstruction at the level similar to *in-vivo* experiments. The additive Gaussian noise was applied to all sinogram volumes to simulate the electronic and thermal noise. Finally, simulated sinograms were reconstructed by maximum likelihood expectation maximisation (MLEM) algorithm by using Tomographic Iterative GPU-based Reconstruction Toolbox (TIGER) (Ander Biguri 2016). Each reconstructed frame was smoothed by using 10mm FWHM Gaussian kernel.

#### Quantitative assessment metrics

To quantitatively compare the activation maps obtained by GLM and ICA in simulation study. Two metrics, sensitivity and specificity, are defined as below

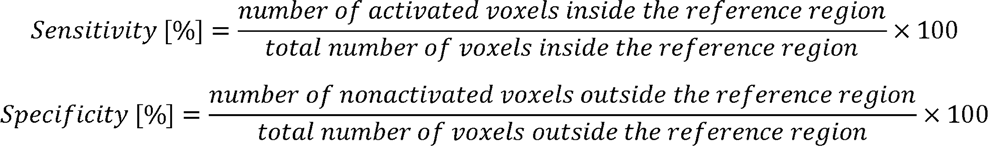

The sensitivity measures the percentage of correctly identified activated voxels. The specificity measures the percentage of nonactivated voxels that have not been wrongly activated.

These metrics were used to compare binary activation maps (*t*-score map in GLM and *z*-score map in ICA) insides the brain. However, we cannot directly compare *t*-score and *z*-score as they possess different statistical meaning. In order to compare them, we threshold and binarize the activation maps based on their statistical significance levels. In the GLM activation maps, sensitivity and specificity are calculated for voxels with positive *t*-scores with significant level *p* < 0.1 (FDR adjusted). In ICA, we set threshold *z* ≥ 1.6 corresponding to approximately *p* < 0.1 for two-tailed normally distributed *z*-scores.

### 2.4 *In-vivo* fPET Experiments

All experiments involving human subjects have been reviewed and approved by the Monash University Human Research Ethics Committee, in accordance with Australian Code for the Responsible Conduct of Research (2007) and the Australian National Statement on Ethical Conduct in Human Research (2007).

Data and code from this study are available on request from the corresponding author pending the institute Ethics approval.

#### Task-fPET experiments

Datasets from six healthy subjects (24.3±3.8 yr, 5 females) were used to compare GLM and fPET in a visual task-fPET study. Subjects were scanned using a 3T Siemens Biograph mMR (Siemens Healthiness, Erlangen, Germany) MR-PET scanner. The full cohort dataset was reported in our previous work in Jamadar et al (2019).

All subjects were asked to take part in three visual stimulation tasks during scan; the experimental design was adapted from Villien et al., 2014. Each subject was infused with 100MBq (administered dose 95.8±5.9 MBq) FDG at a constant rate (36mL/hr) over 90 minutes. All subjects were instructed to lie in quietly under resting condition with eye closed in the first 20 minutes to allow the sufficient FDG accumulation in the brain, and a series of non-functional fMRI scans were conducted during this period. A 10-min full checkerboard stimulation started at 20 minutes after subjects were instructed to open eyes. The checkerboard was shown on screen for 120-sec, and then 32-sec on and 16-sec off design was implemented. During the ‘on’ periods, the visual stimulus was a circular checkerboard of size 39cm (visual angle 9°) presented on a black background. The checkerboard flickered (i.e., fields alternated black/white) at 8Hz. During the ‘off’ periods, subjects rested eyes open while viewing a white fixation cross of size 3cm (visual angle 0° 45’) presented on a black background. After the first full-checkerboard stimulation, there were 15-min eyes-closed resting period followed by two 5-min half hemifield stimulation with 5-min eyes-closed resting period in-betweens. Then, there was another 20-min eyes-closed resting period before the second 10-min full checkerboard stimulation. The PET data acquired during the first full-checkerboard were used to validate the GLM and ICA methods. There were thirty minutes PET data for each subject, including 10 mins resting before the stimulation, 10 mins full-checkboard stimulation followed by another 10mins of resting.

The PET list-mode data were binned into 30 frames (1 min per frame). The pseudo-CT attenuation map was used to correct the attenuation for all acquired data (Burgos et al. 2014). Ordinary Poisson-Ordered Subset Expectation Maximization (OP-OSEM) (3 iterations, 21 subsets) algorithm with Point Spread Function (PSF) correction was used to reconstruct 3D volumes with matrix size 344×344×127 (Voxels size: 2.09×2.09×2.03 mm^3^). A guided motion correction method using simultaneously acquired MR images was applied to the 4D PET volumes for each subject (Chen et al. 2019). A 12-mm FWHM Gaussian post-filtering was applied to each 3D volume. The PET images for each subject were registered to the 2mm MNI-152 template.

#### Resting state fPET

For the resting state fPET study, healthy adults (n=28, 19.6±1.7 yr, 21 female) underwent simultaneous BOLD-fMRI/FDG-PET on a 3T Siemens Biograph mMR. All subjects were instructed to lie quietly at rest with their eyes open. Each subject was infused with 260MBq (actual administered dose 234±19 MBq) FDG at a constant rate (36mL/hr) over 95 minutes. Several MRI acquisitions were performed during the PET scan; UTE, T1 MPRAGE, T2-SPC, pASL, and SWI images were acquired in the first 30-mins; followed by 6 consecutive 10-min T2* EPI (TR=2450ms, TE=30ms, FA=90°, 64×64×44, resolution=3×3×3mm^3^) acquisitions. Here, we report the results of the FDG-PET. The PET list-mode data were binned into 60 frames (1 min per frame) aligned to the 6 EPI sessions.

The reconstruction and pre-processing procedures were identical to the task fPET experiment.

## 3. Results

### 3.1 Simulation Experiments

#### Simulated fPET images

A total of 90 frames of PET images were simulated and the length of each frame was 1min. The first 10 frames were discarded due to the extremely low FDG counts. The simulated fPET images based on the kinetic model were shown in Figure 3. Four image volumes are shown at 30 min, 45 min, 60 min and 80 min. Their equivalent FDG activities were 0.34 MBq, 0.64 MBq, 1.00 MBq and 1.56 MBq during the one minute binning window, respectively. The simulation demonstrated the increased FDG activity over time as shown in Figure 2a.

**Figure 3:**
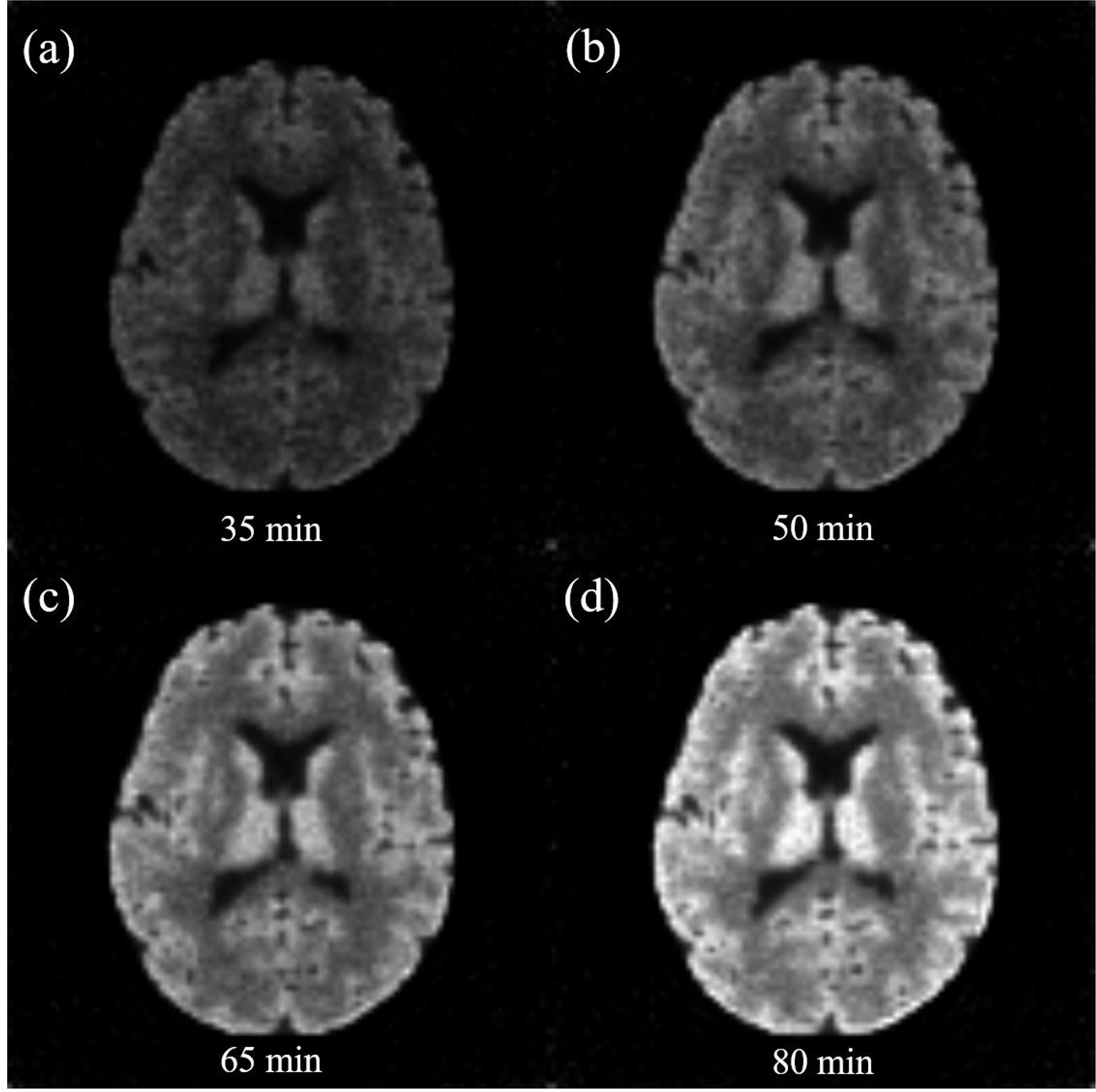
Simulated fPET images reconstructed using the TIGER toolbox. The frame length is 1 min for all image volumes, consistent with our in vivo data. **(a):** image at 35min with FDG activity 0.34MBq, **(b):** image at 50min with FDG activity 0.64MBq, **(c):** image at 65min with FDG activity 1.00MBq, **(d):** image at 80min with FDG activity 1.56MBq. It demonstrates increased activity signals over time.

#### Comparison of GLM and ICA on simulated fPET datasets

Subject level GLM and ICA were performed on the simulated fPET data. To evaluate the performance of GLM, we also used the ground truth regressors for each task, which were given in Figure 2c and 2d. For ICA, the number of independent components was set to five. Two activation maps obtained from the first component were associated with the two designed tasks.

The results of the visual task using the GLM with the ramp regressor, GLM with ground truth regressor, and ICA are shown in Figure 4. Overall, all methods identified the correct activation in the virtual cortex (Figure 4a-4c). The curve fitting results in Figure 4d confirmed that the FDG activity signal (solid line) can be reasonably fitted by the ramp regressor (dash line) with a small degree of error, however the errors become greater at the signals after 70 mins. When the ground truth regressor was applied, and the fit was nearly perfect (Figure 4e). The corresponding activation maps in Figure 4b also provided higher *t*-score than Figure 4a (all *p* < 0.1, FDR adjusted) compared between the two GLMs. The ICA timecourse in Figure 4f represents a relatively signal change related to the task, and correctly identified the increased FDG time activity during two task stimulation periods from 20min to 30min, and 70min to 80min.

**Figure 4:**
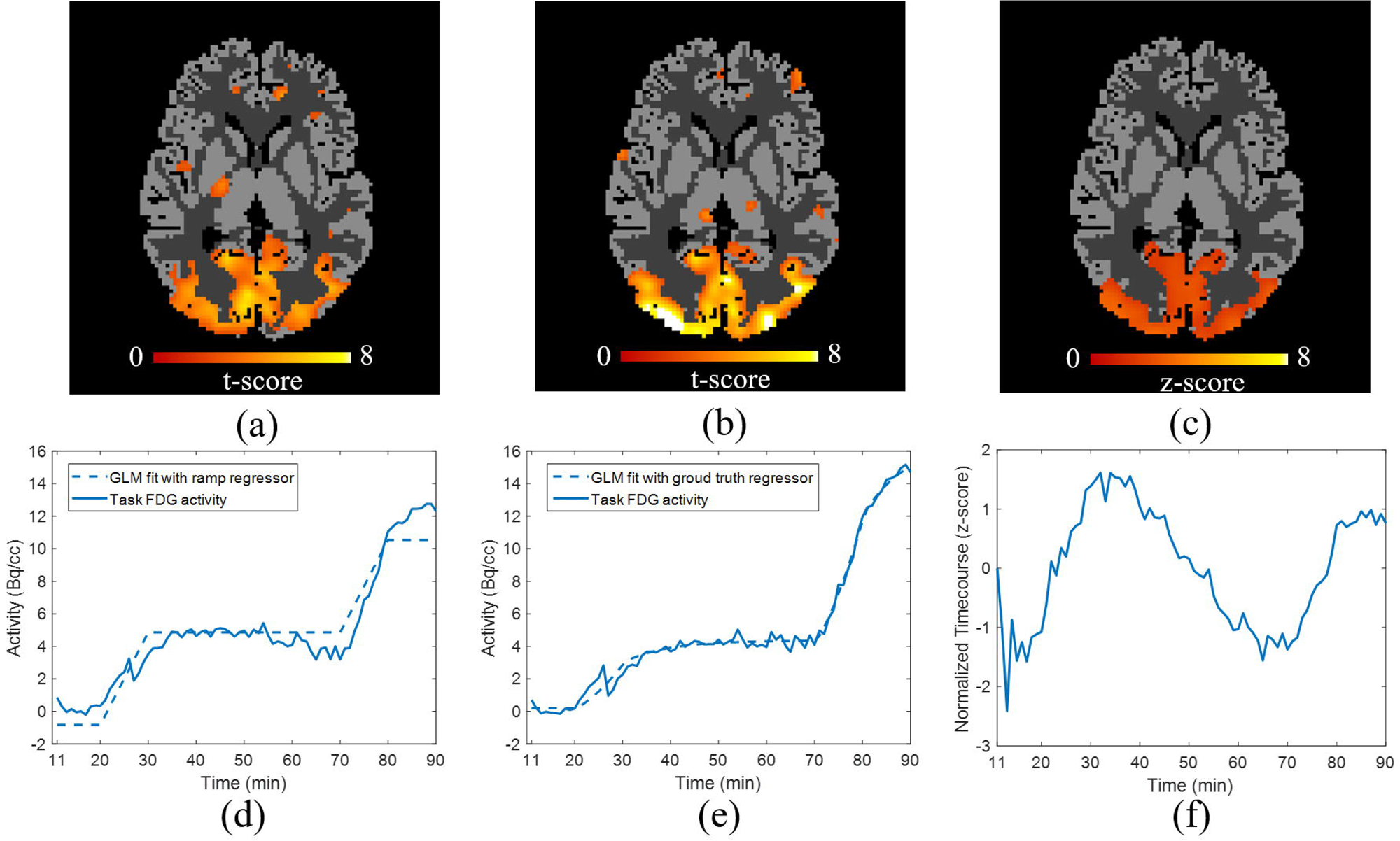
Comparison of the visual task results from the simulation experiment. **(a):** activation map of GLM using the ramp regressor (p<0.1, FDR adjusted), **(b):** activation map of GLM using ground truth task regressor (p<0.1, FDR adjusted), **(c):** activation map of ICA (z>1.6), **(d):** GLM fitting using the ramp task regressor, **(e):** GLM fitting using the ground truth regressor, **(f)**: ICA timecourse (z-scored).

The motor task results for the two GLMs and ICA are shown in Figure 5. All three activation maps in Figures 5a, 5b and 5c demonstrated consistent activation in the motor region. The actual fitted results (Figures 5c-5e) further confirmed our observation from the visual task experiments.

**Figure 5:**
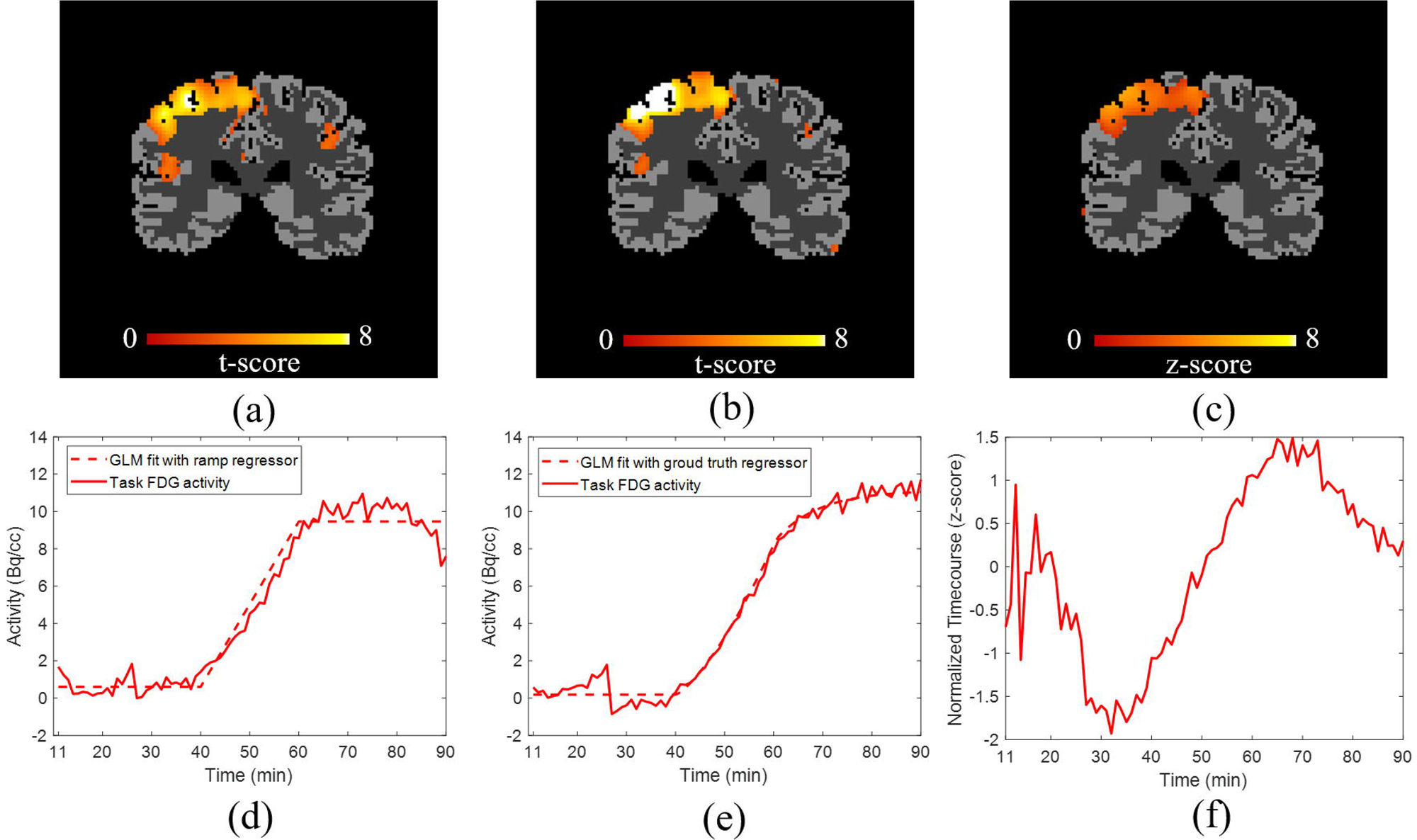
Comparison of the motor task results from the simulation experiment. **(a):** activation map of GLM using the ramp regressor (p<0.1, FDR adjusted), **(b):** activation map of GLM using the ground truth regressor (p<0.1, FDR adjusted), **(c):** activation map of ICA, **(d):** GLM fitting using the ramp regressor, **(e):** GLM fitting using the ground truth regressor, **(f)**: ICA timecourse (z-scored).

The sensitivity and specificity of activation maps are provided in Table II. Both visual and motor tasks were evaluated. In general, GLM provided better sensitivity than ICA in both experiments, but with lower specificity. This suggests that GLM can more accurately identify the regions of interest, but with higher number of false activations outside the region of interests. By using the ground truth regressors in GLM, we observed noticeable increased sensitivity in the visual task experiment. Compared between the two tasks, visual task had lower overall sensitivity attributable to its more complex underlying signal shape. All methods show specificity above 90% suggesting accurate identification of task related signal changes.

**Table II:** Comparison of sensitivity and specificity for GLM using ramp regressor, GLM using ground truth regressor and ICA from the simulated visual and motor task experiments.

|  | Sensitivity |  | Specificity |  |
| --- | --- | --- | --- | --- |
|  | Motor task | Visual task | Motor task | Visual task |
| GLM using ramp regressor | 96.9% | 79.6% | 96.5% | 93.3% |
| GLM using ground truth regressor | 97.5% | 86.0% | 95.6% | 92.4% |
| ICA | 95.8% | 76.7% | 98.3% | 97.2% |

### 3.2 *In-vivo* fPET Experiments

#### Task-fPET experiment

Both GLM and ICA methods were applied on the *in-vivo* visual task experiment. We performed both subject level and group level analyses. During the PCA pre-processing for ICA, the number of principle components for each subject was set to twenty, and was further reduced to ten during the group level stage. The component related to the visual system was selected.

The group level activation maps are shown in Figures 6 using the GLM and ICA methods, respectively. Overall both methods successfully identified activation in the visual cortex (Figures 6a and 6c), but the extent of the activations were slightly different across the methods by visual comparison. Using ICA, an additional negative brain activation in the superior sagittal and straight sinuses was also given in Figure 6d. The fitting curve in Figure 6b demonstrated an accurate GLM fit at the group level. Similarly, the ICA timecourse in Figure 6e also identified increased relative FDG activity uptake (red) during the task stimulation and reduced relative uptake (blue) in the negative activation region. Both methods demonstrated the clear activation in occipital lobe at group level though they were not perfectly overlapped. At subject level, we found consistent ICA activations in five out of six individuals, and only four out of six individuals for GLM. The results of each individual subject are provided in Supplementary materials I.

**Figure 6:**
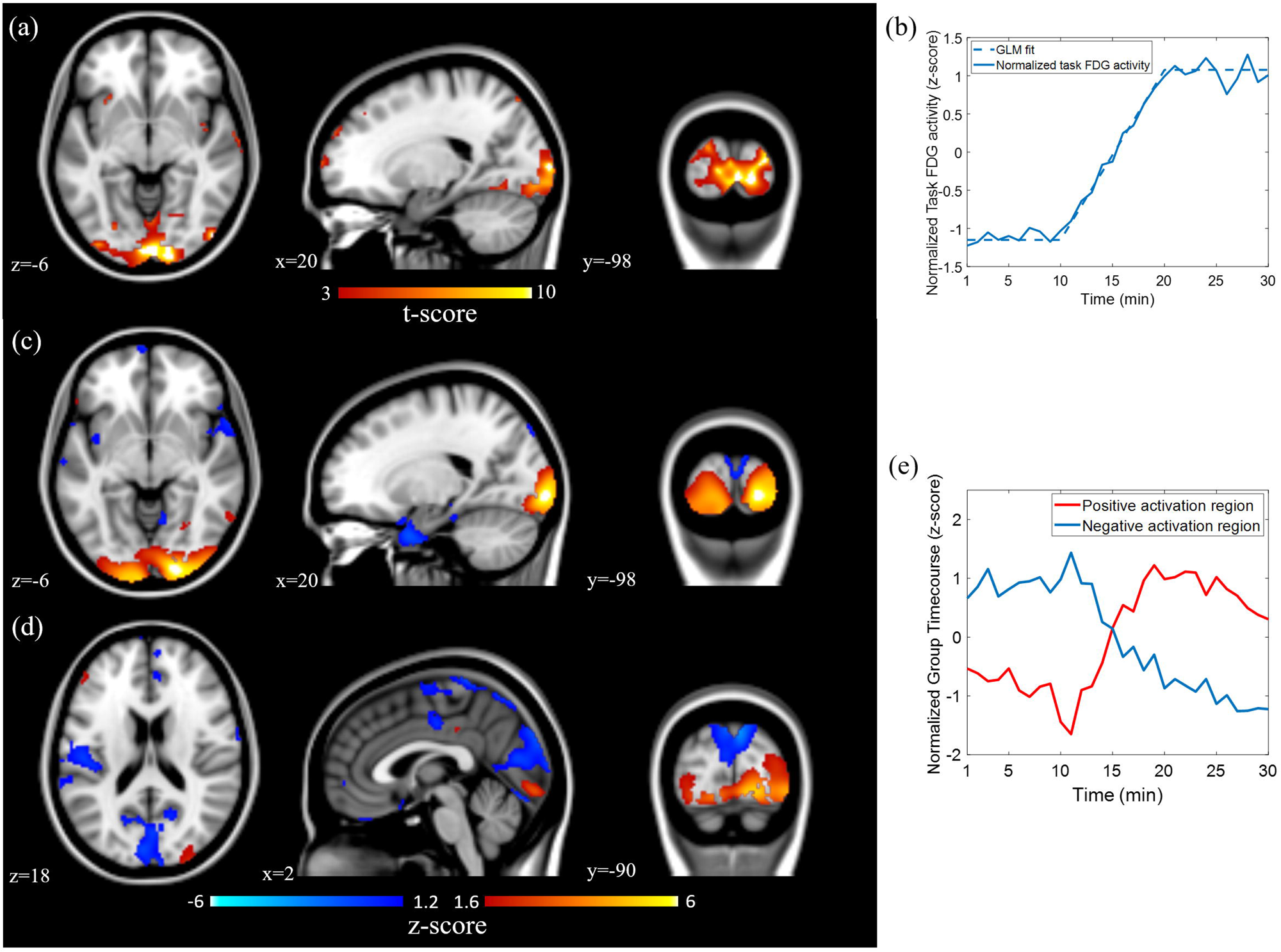
In-vivo visual experiment results. **(a):** orthogonal view of GLM activation map (t-score >3). **(b):** normalized task fitting curve corresponding to the activation map. **(c):** orthogonal view of ICA activation map in visual cortex. **(d):** the negative ICA activation map shows activation in sagittal and straight sinuses. **(e):** normalized group timecourse (z-scored) corresponding to the activation map.

#### Resting state fPET experiment

As GLM requires a functional regressor that is absent in resting state paradigms, it is ill-suited to resting state functional analysis. Thus, only ICA was investigated. The data dimension of each subject was reduced to 40 using PCA, and then normalized to remove the global baseline. The data dimension of concatenated group data was further reduced to 40. A total of 20 activation maps were obtained using the FastICA toolbox. Figure 7 presents the orthogonal view of all components. Overall, there were two networks located in the frontal lobe (Figure 7a and 7b), three located in the parietal lobe (Figure 7c-7e), five in occipital lobe (Figure 7f-7j), two in the sub-cortical regions (Figure 7k and 7l), one in cerebellum (Figure 7m), one in mesial frontal/parietal (Figure 7n), one in frontal/temporal (Figure 7o) and one in frontal/parietal (Figure 7p). Figure 7q-7t were labelled as noise or mixed signal/noise components. Detailed axial views of these activation maps were provided in Supplementary materials II. Several networks resemble fMRI resting state networks, such as default mode network Figure 7c, primary visual Figure 7f and Figure 7j, secondary visual Figure 7g and cerebellum Figure 7m, but others cannot be easily mapped to canonical fMRI resting state networks.

**Figure 7:**
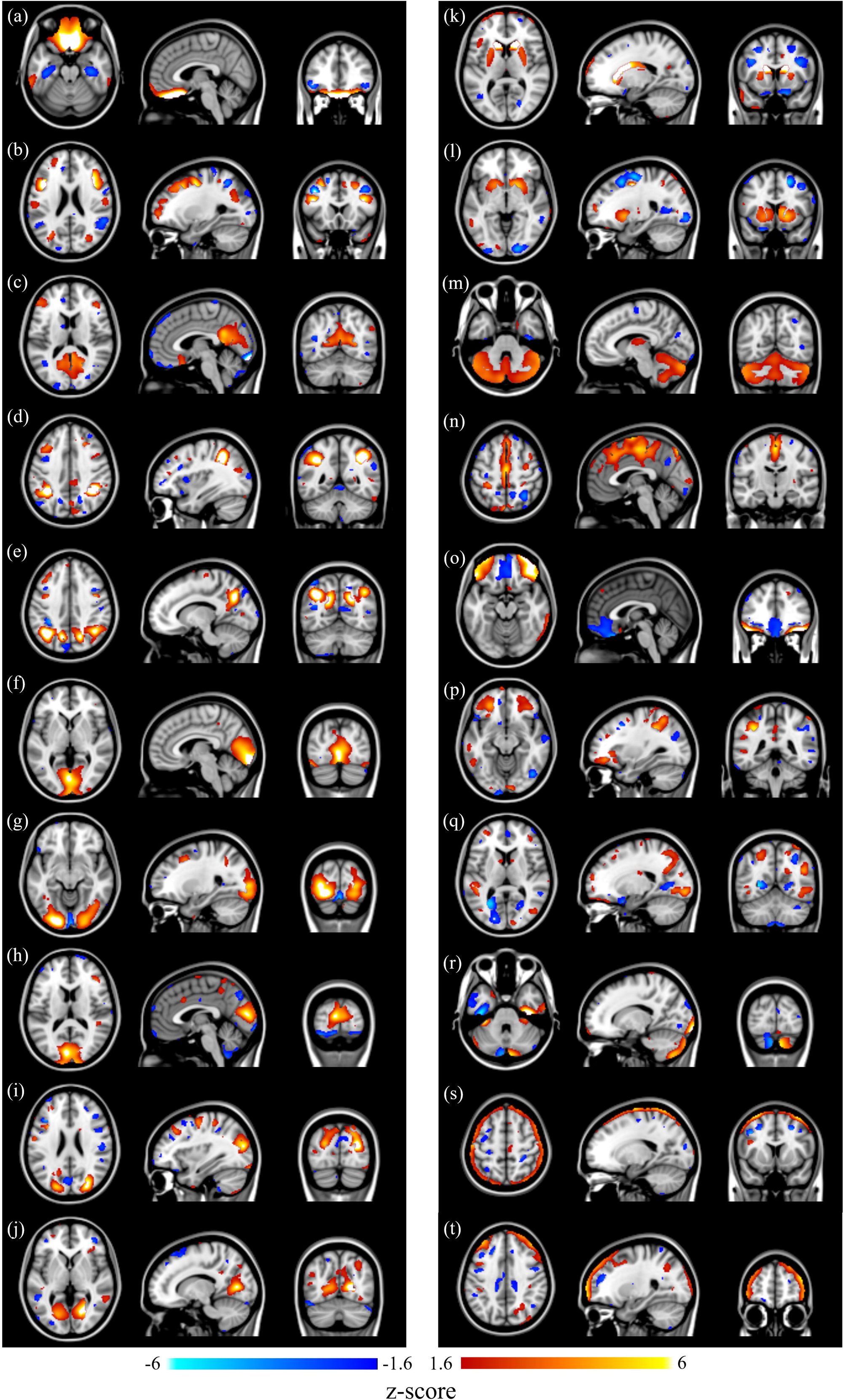
fPET resting state brain networks. **(a)-(b)** present two networks in frontal lobe. **(c)-(e)** present three networks in parietal lobe. **(f)-(j)** present five networks in occipital lobe. **(k)-(l)** presents two networks around ventricle. **(m)** cerebellum. **(n)** mesial frontal/parietal. **(o)** frontal/temporal. **(p)** frontal/parietal. **(q)-(t)** noise or mixed signal/noise.

Furthermore, within several resting state metabolic networks, we also observed widespread anti-correlated brain regions. These anti-correlated brain regions are shown as positive (red) and negative (blue) brain activations (in Figure 7), respectively.

## 4. Discussion

In this work, we investigated two types of fPET analysis methods including a model-based GLM method and a data-driven ICA method. We show that both methods can successfully identify task-related FDG activations using both *in-silico* and *in-vivo* task-fPET experiments. The GLM demonstrates higher sensitivity than ICA probably attributable to the *a priori* knowledge used during data fitting, whereas the ICA shows slightly better specificity in the simulation study. Additionally, we found that ICA produced more consistent activations across individuals, although a larger investigation is required to validate and quantify this discrepancy. In the resting state fPET analysis, the data driven ICA method was able to derive brain resting state metabolic networks.

Unlike fMRI, the analysis of fPET requires an accurate estimation of the cerebral metabolic baseline, especially when using GLM. The metabolic baseline accounts for over 90% of the total time activity signals; the task-related time activity is only a small percentage of the signal. Any residuals in the baseline estimation can introduce bias when fitting the task design matrix. Hahn et al. (2016) applied a 3rd order polynomial to estimate a basal metabolic baseline regressor and demonstrated that the metabolic baseline is brain region dependent. Compared with GLM, ICA avoids the explicit model-fit for task stimuli, and the baseline removal in ICA is simply to remove global signals and maintain regional metabolic variations. As a result, ICA reduces false positive activation compared with GLM, as demonstrated in our simulation study.

The GLM analysis of fPET requires an *a priori* knowledge for the task regressor. A ramp function is approximated as the task regressor which has produced reasonably accurate activation maps. However, as shown in the simulation study, the ground truth regressor actually takes a more complex shape than a ramp function since the FDG activity change is an integration effect from the changes in the brain kinetics. This effect was clearly demonstrated in the simulation study. The task-only time activity signal increases beyond the duration of the task stimulation. Therefore, additional investigation into regressor design is warranted, and care should be taken when inferring GLM fitting results in neuroscience applications.

Compared with fMRI ICA, there are several distinct features in the fPET ICA methods. First, the need to normalize data to remove a global baseline as well as to reduce inter-subject viability during the group level analysis. Second, widespread anti-correlated brain regions within a single independent component are identified using the ICA method. These anti-correlated regions can potentially lead to interesting findings when evaluating their functional metabolic connectivity, however they may be spurious manifestations of the global baseline removal, as has been observed in fMRI following global signal regression (Murphy and Fox 2017). Similar to the application of ICA in fMRI data analysis, the number of independent components is empirically determined (Du and Fan 2013; Hui et al. 2011). However, some efforts have been made to mitigate this empirical approach by using minimum description length (MDL) (Calhoun et al. 2001b; Li, Adali, and Calhoun 2007). These methods may improve the performance of ICA for fPET analysis.

In this work, a template image and a tissue kinetic model were used to simulate PET sinogram data and a thinning Poisson process was used to generate the dynamic PET data. The thinning Poisson process has been used by others for the evaluation of PET image reconstruction methods (Kim et al. 2018). However, the simulated data would be more realistic if a full Monte Carlo simulation of positron emission decay and the PET detection process at the coincidence event level (e.g. using the GATE software (Jan et al. 2004)) was used.

One current limitation of fPET is that the temporal resolution of fPET time activity is significantly lower than BOLD fMRI. The temporal resolution of task fPET has been recently improved from 1 min (Hahn et al. 2016; Villien et al. 2014) to the below 20 seconds (Jamadar, Ward, Carey, et al. 2019; Rischka et al. 2018). However, this is still much slower than the seconds or even sub-seconds temporal resolution in BOLD fMRI. It is also worthwhile to note that each fPET frame measures the mean FDG activity within the frame duration. Only a small fraction of this activity is likely to be due to recent stimulated or spontaneous neural responses compared to the integral of all past activity.

Conventional dynamic PET signals can be modelled using kinetic compartment models (Phelps et al. 1979; Sokoloff et al. 1977). The determination of kinetic rate constants could potentially be used to analyze variation in fPET FDG uptake across subjects when presented with external stimuli. However, the development of parametric analysis methods for fPET signal uptake is beyond the scope of this manuscript. Careful design and optimization of the fPET analysis method will be a critical step in future studies for interpretation of the fPET findings.

FDG PET imaging has long been a proxy for investigating brain metabolism (Passow et al. 2015; Tomasi et al. 2017). fPET, extended from the conventional static PET, can potentially be useful in the investigation of the complex hemodynamic and metabolic relationship in simultaneous MRI and PET study (Wehrl et al. 2013), resting state functional metabolic covariation (Amend et al. 2019; Di, Biswal, and Alzheimer’s Disease Neuroimaging 2012), and application in neurodegeneration studies by assessing metabolic networks (Eidelberg 2009). With efforts invested into fPET acquisition methods, e.g. the bolus plus continuous infusion, we can further improve its temporal resolution (Jamadar, Ward, Carey, et al. 2019; Rischka et al. 2018). Advanced PET image reconstruction can improve the overall PET image quality (Pamulakanty Sudarshan, Chen, and P. Awate 2018), and these technological improvements can potentially enhance the usefulness of fPET in neuroscience and clinical applications.

## 5. Conclusion

The unique characteristics of fPET signals have been investigated to evaluate analysis methods for task and resting state fPET experiments. Using a simulated task fPET experiment, we quantitatively evaluated the performance of the model based GLM method and the data driven ICA method. Using an *in-vivo* visual task fPET experiment, activation in the visual cortex was identified using both GLM and ICA methods with anatomically similar regions of activation identified. Both methods showed a good agreement in the task fPET experiments, although the GLM showed slightly better sensitivity than ICA. ICA has proven its effectiveness for the resting state fPET analysis. By applying spatial ICA on temporally concatenated fPET datasets, we have presented resting state metabolic brain networks in a cohort of 28 healthy subjects. Overall, fPET provides a unique method to map dynamic changes of glucose uptake in the resting human brain and in response to extrinsic stimulation.

## Acknowledgements

S.Li and Z.Chen are supported by funding from Reignwood Cultural Foundation. S.D. Jamadar is supported by an Australian Research Council Discovery Early Career Research Award (DE150100406). The authors are grateful Richard McIntyre, Alexandra Carey and Jason Bradley for organizing scanning protocol. We thank Edwina Orchard, Irene Graafsma and Disha Sasan in the assistance of data collection; Francesco Sforazzini and Jakub Baran in the assistance of PET data preparation.

