## Supplement I for "Analysis of continuous infusion functional PET (fPET) in the human brain"

Supplementary I

Results of subject level task fPET using GLM and ICA

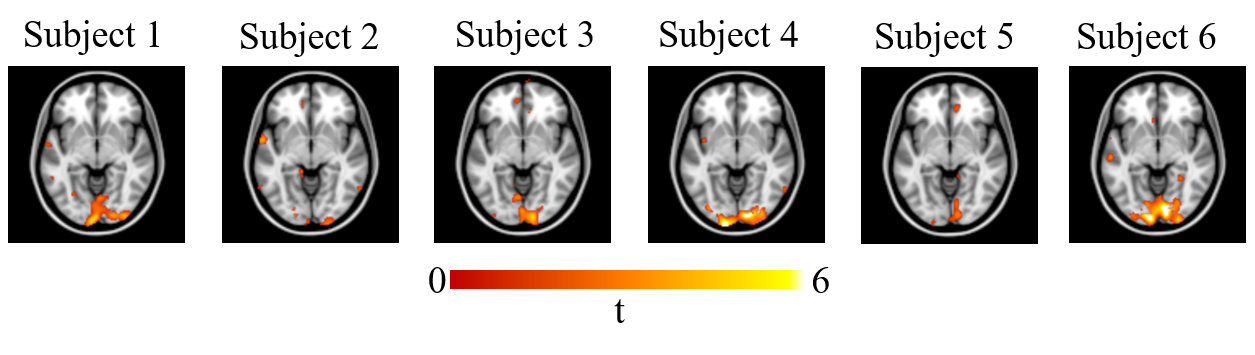

Figure S1 Subject-level GLM activation maps for six subjects (all p<0.1, FDR adjusted)

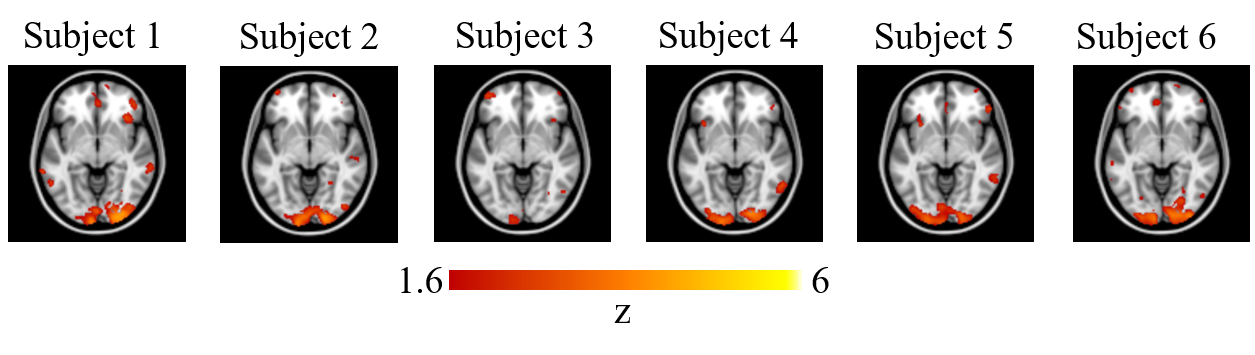

Figure S2 subject-level ICA activation maps back projected from the group results
