## Supplementary figures and images for "Analysis of continuous infusion functional PET (fPET) in the human brain"

### Supplement II

Supplement II

Axial view of resting-state ICA components.

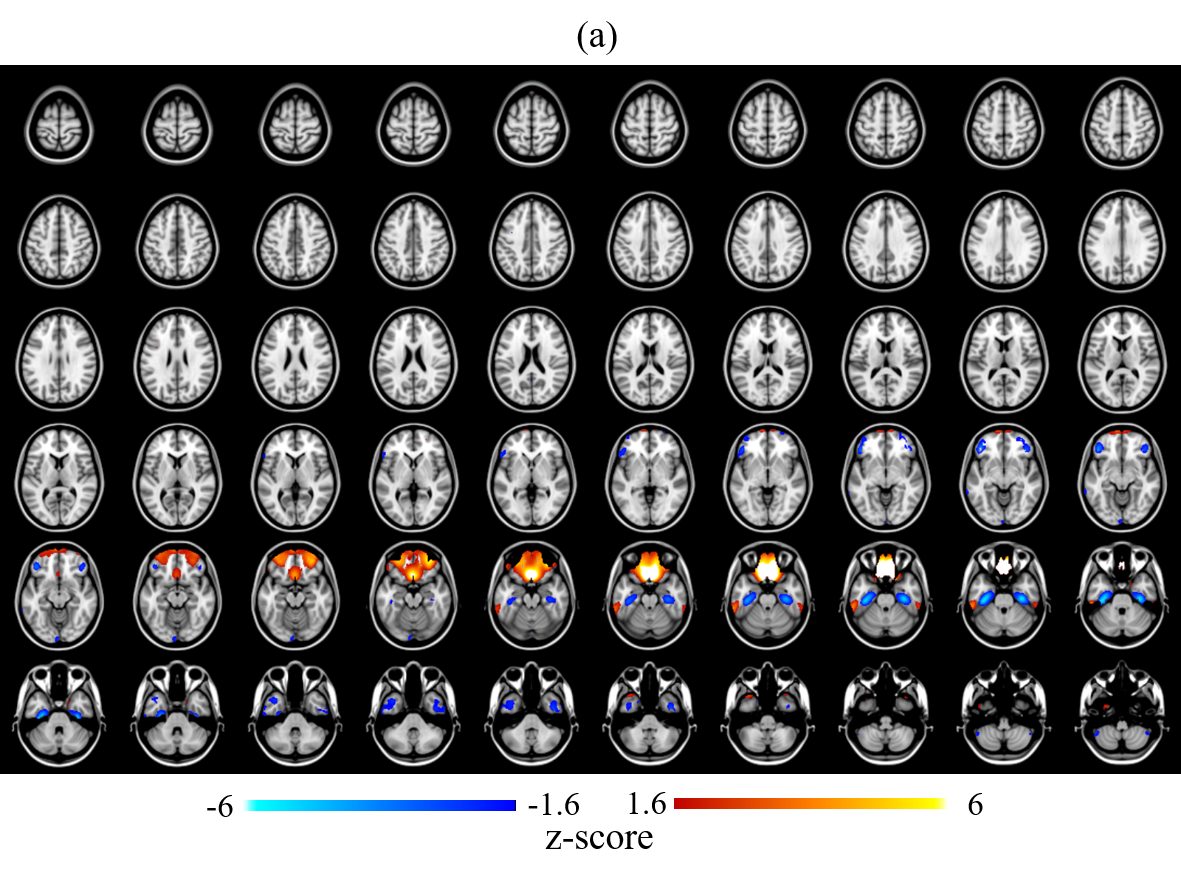

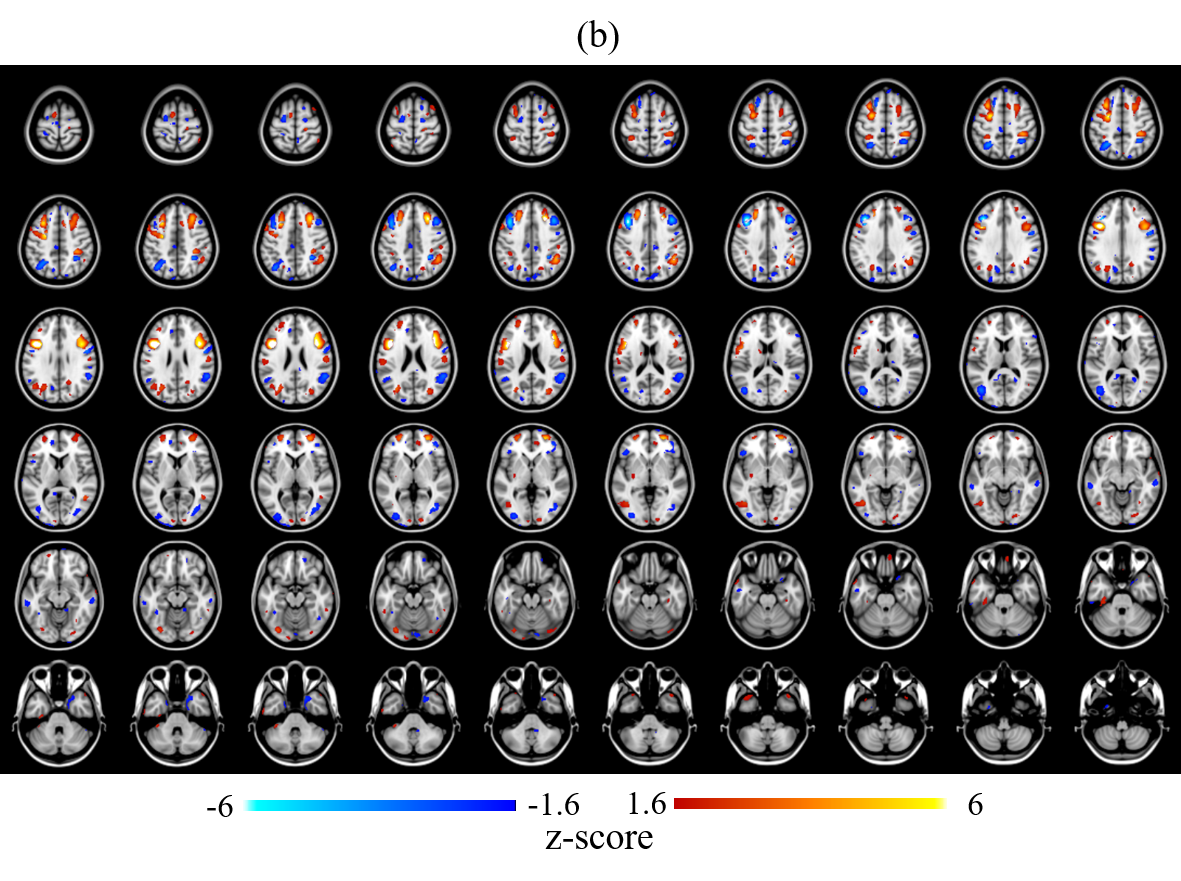

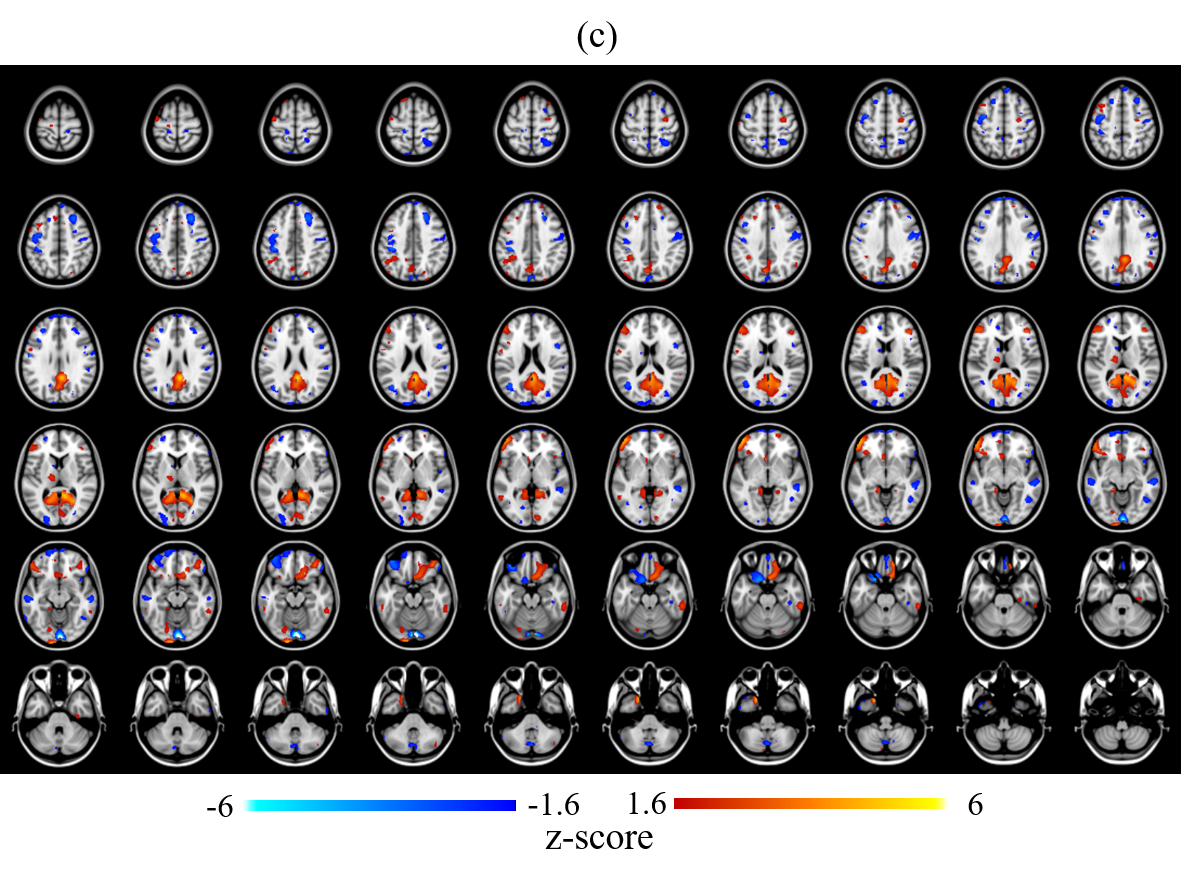

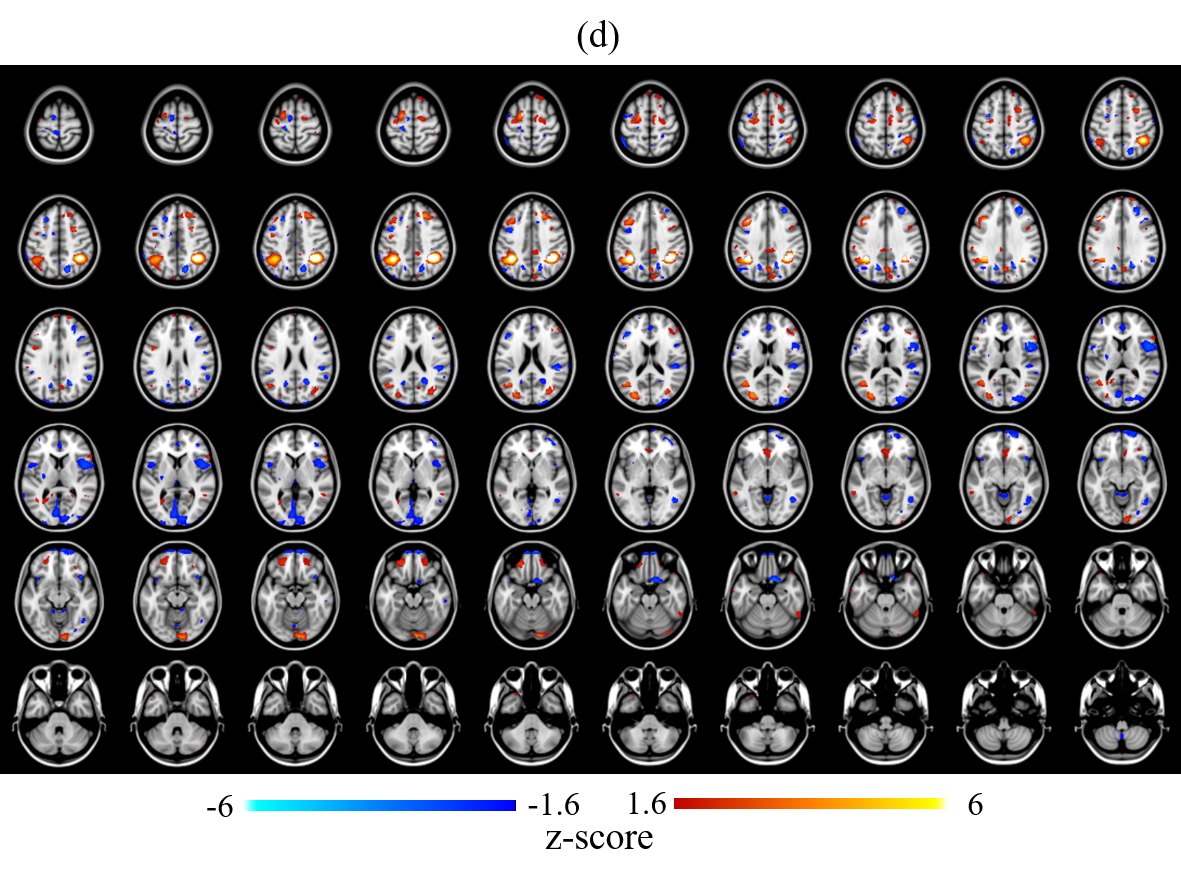

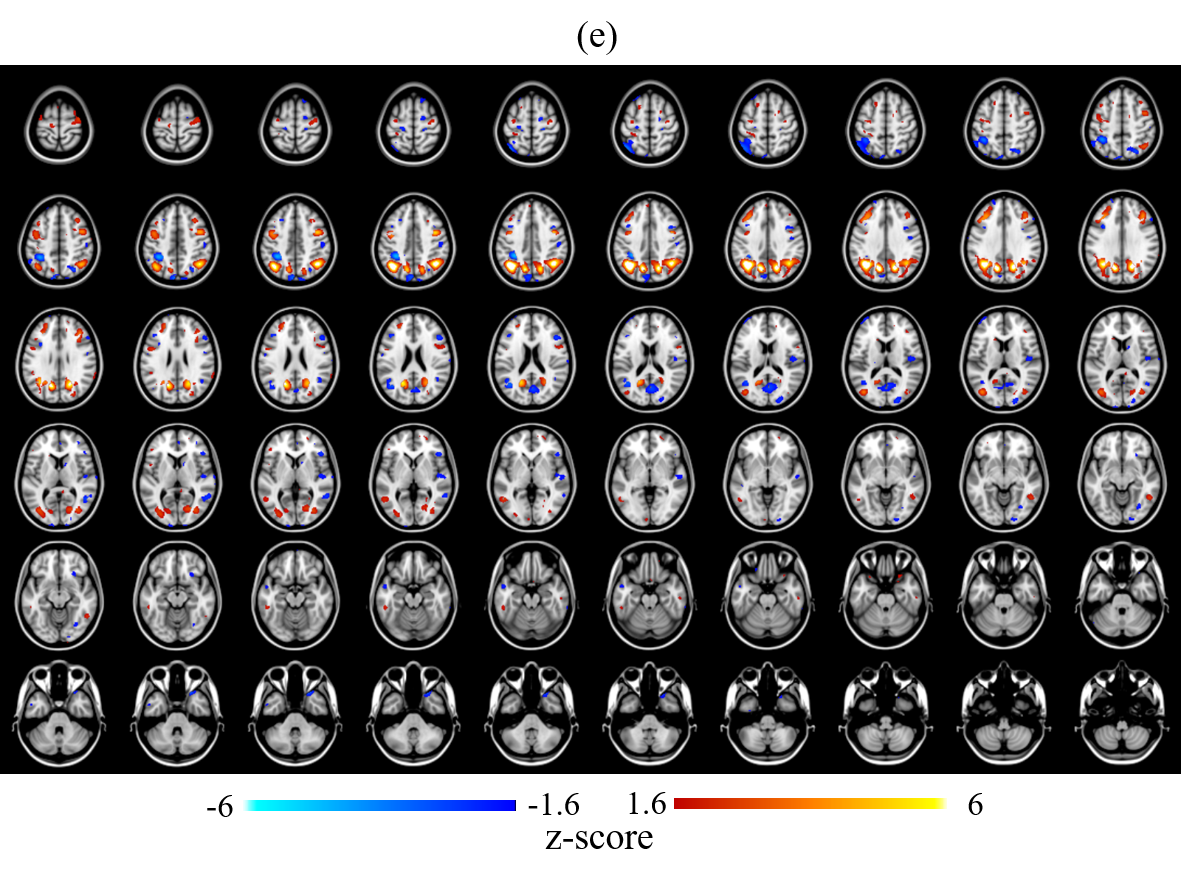

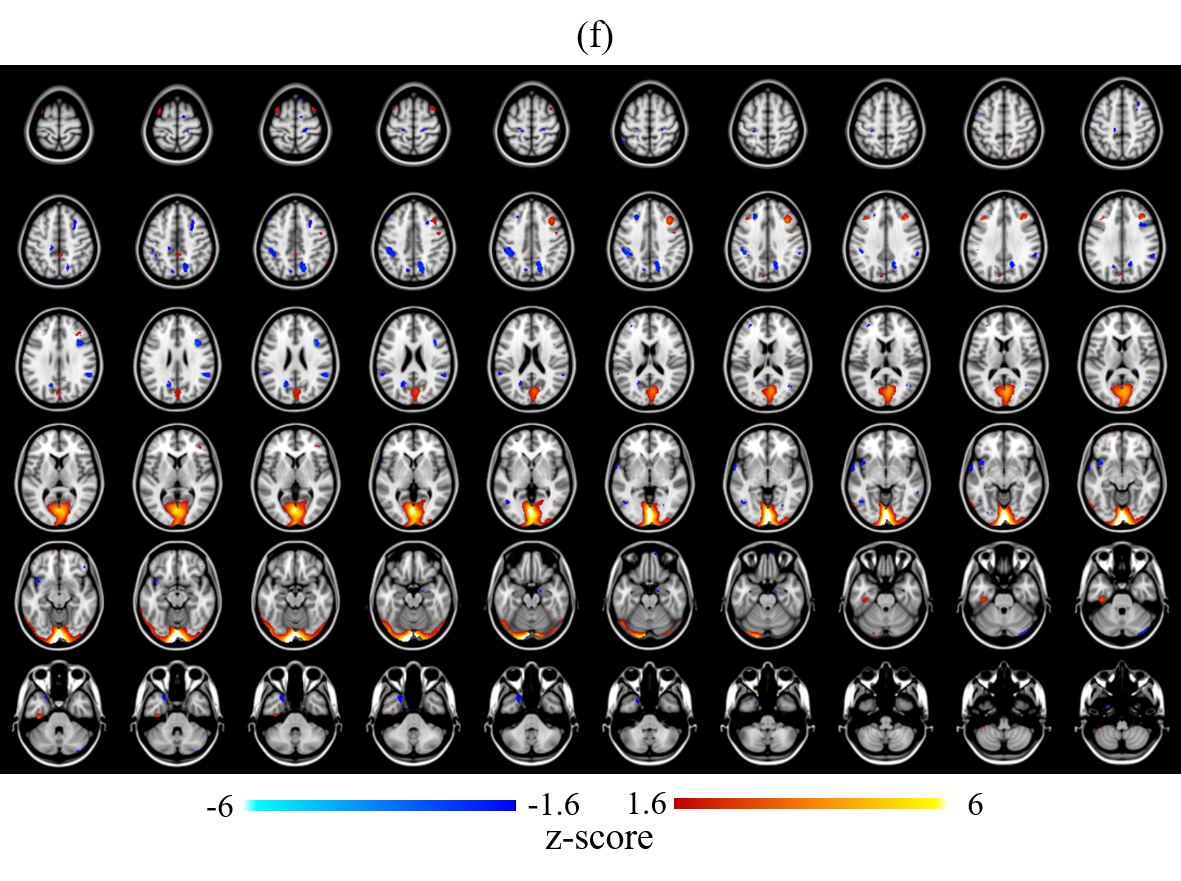

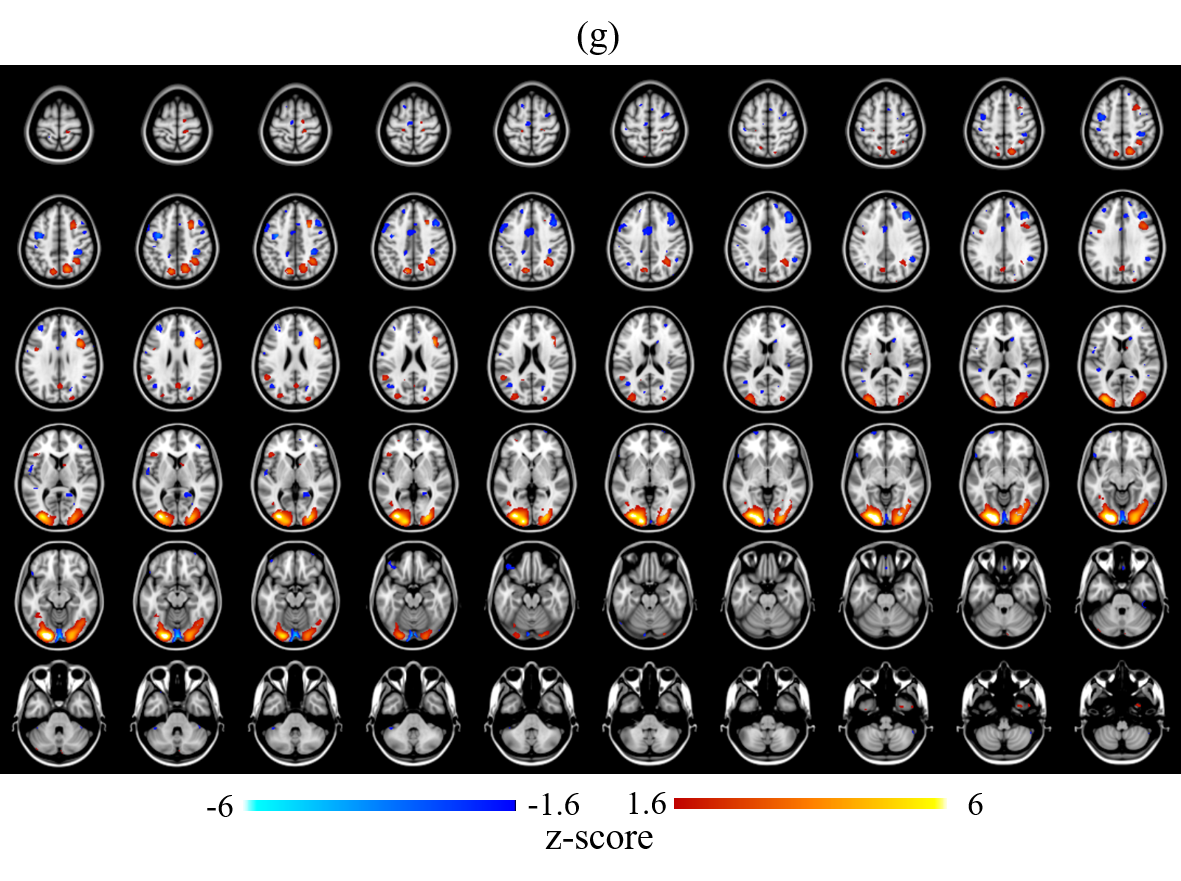

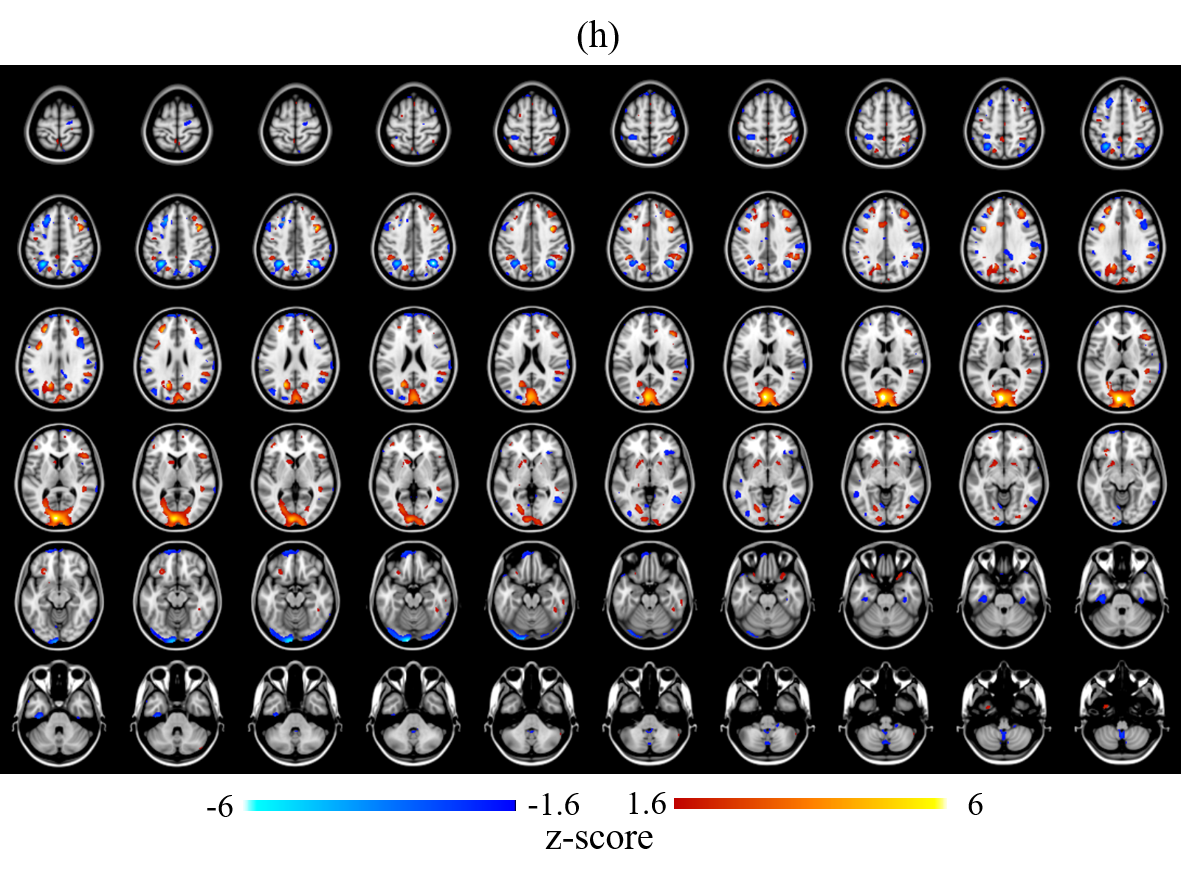

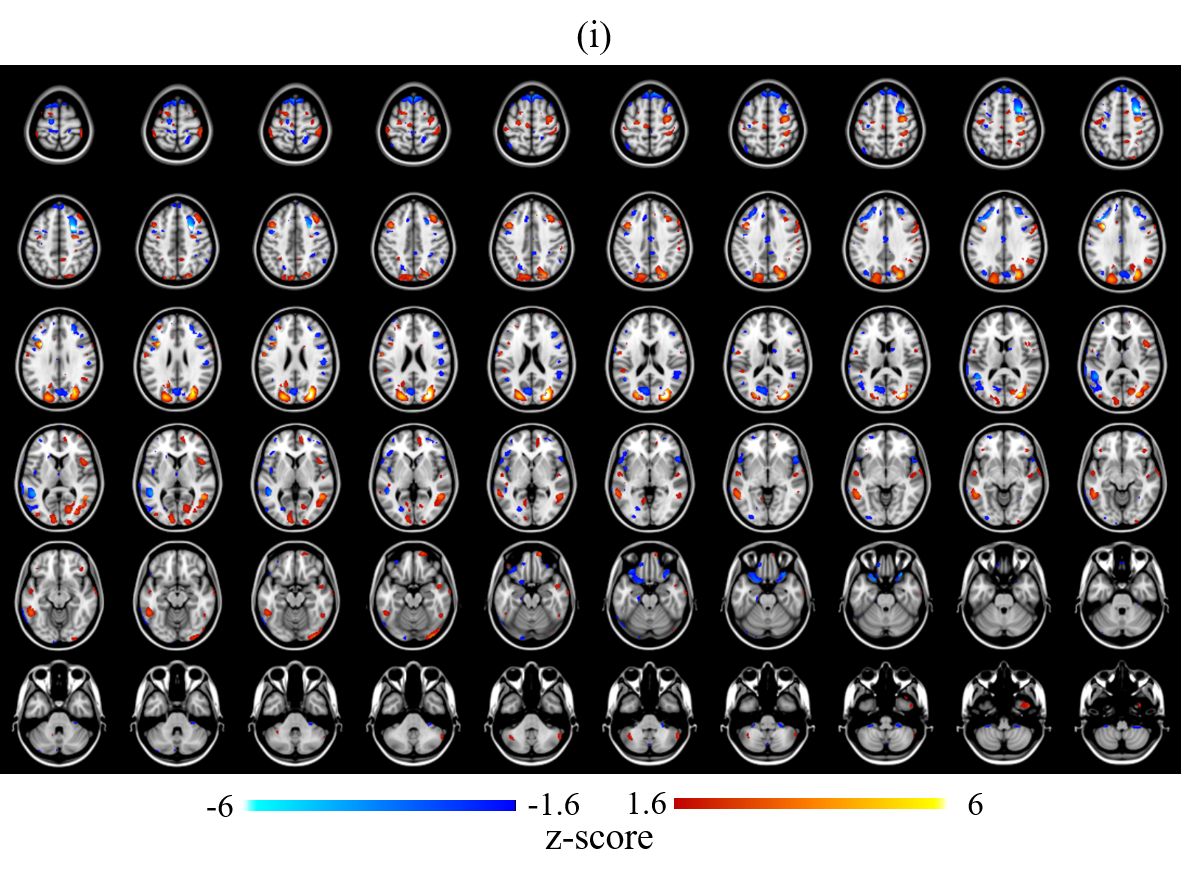

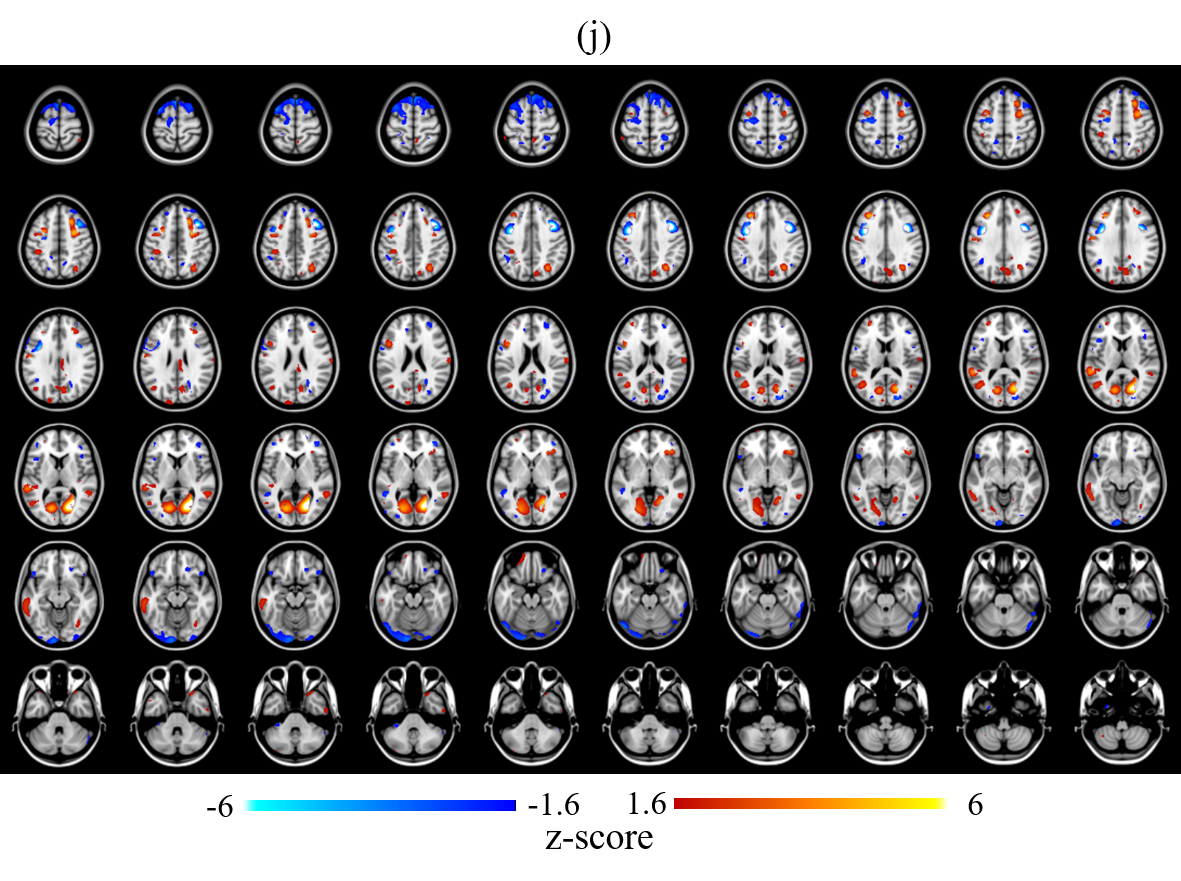

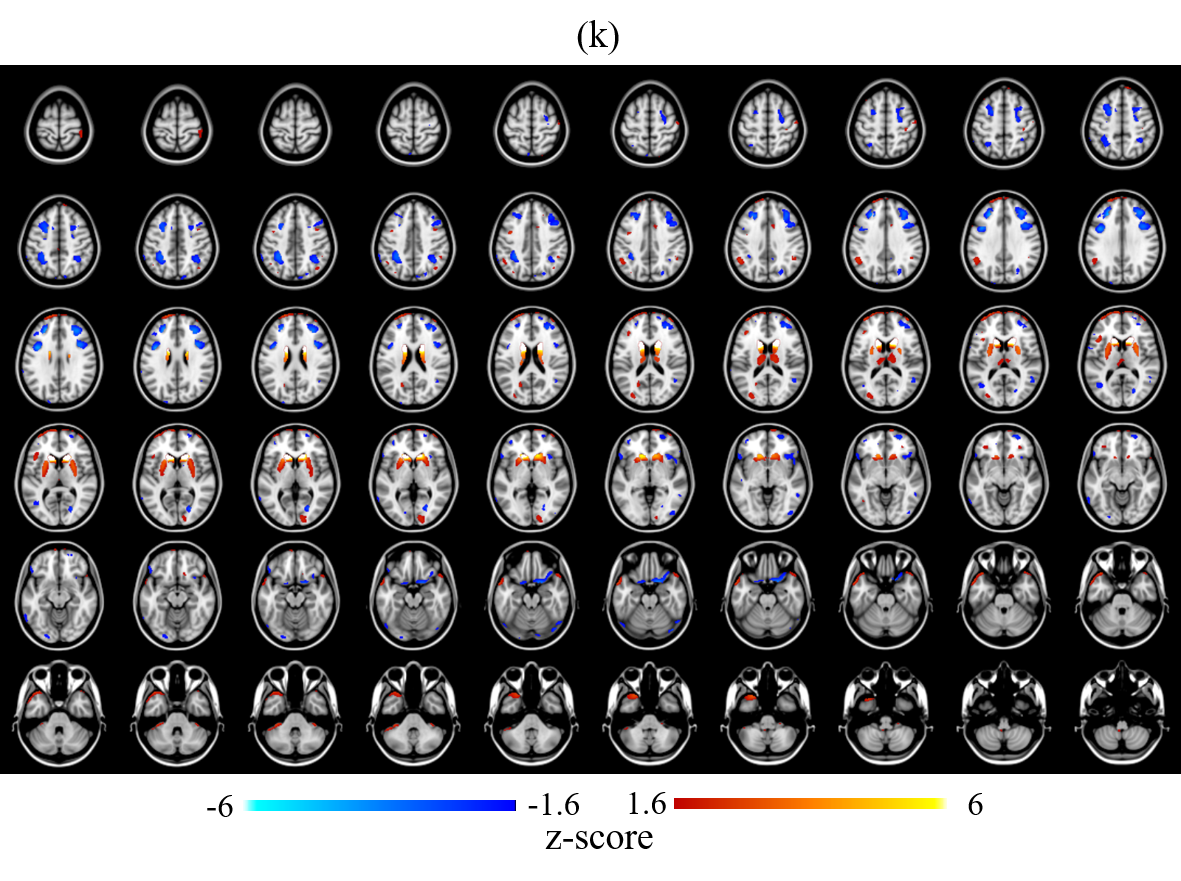

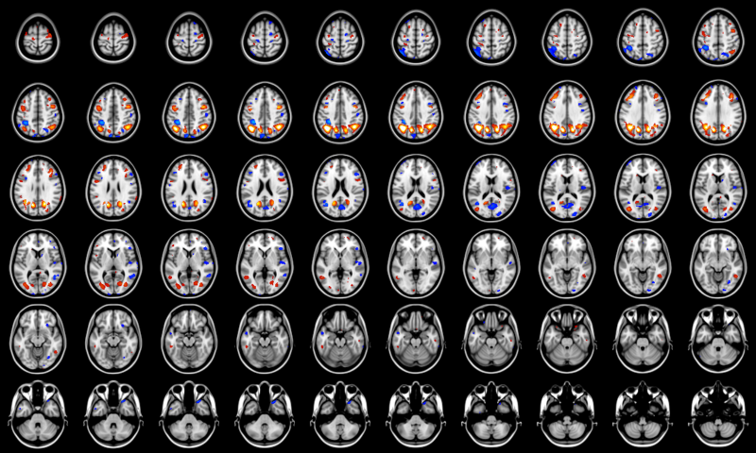

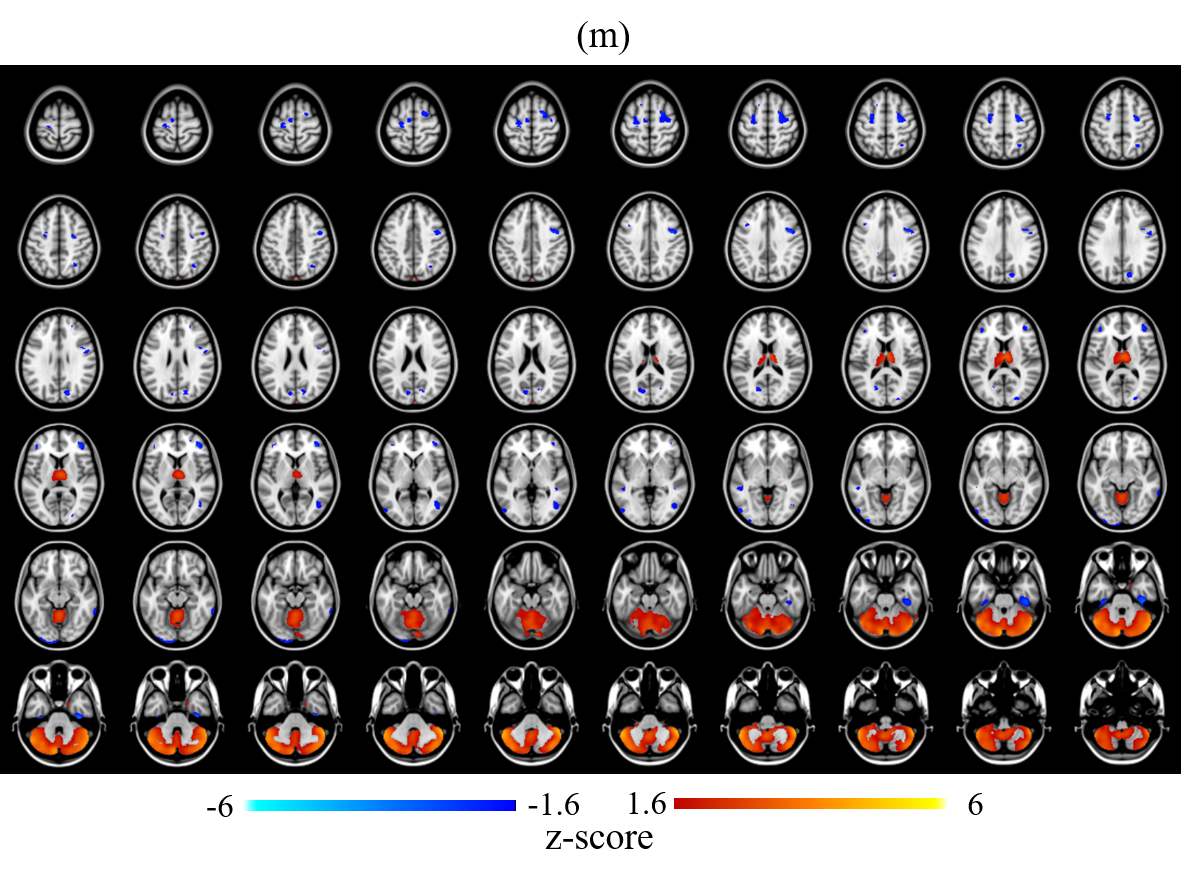

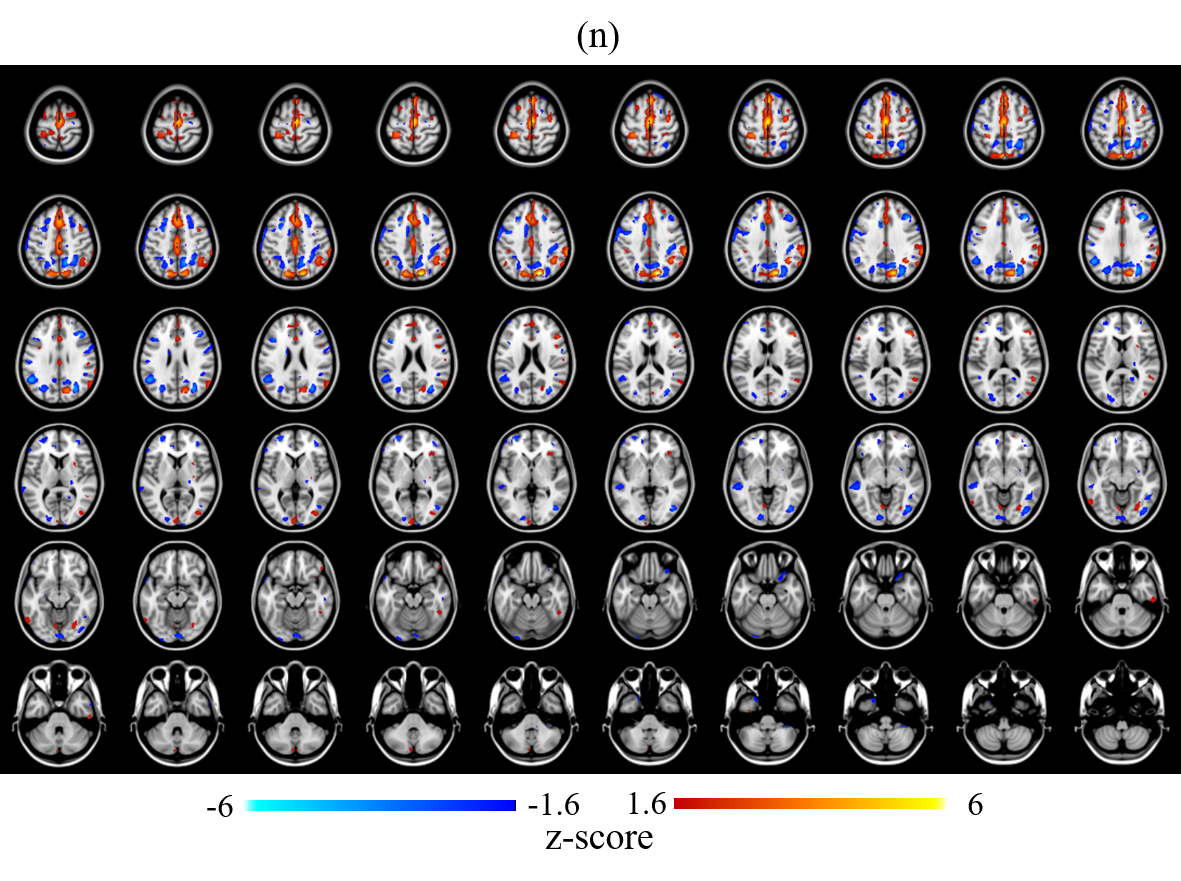

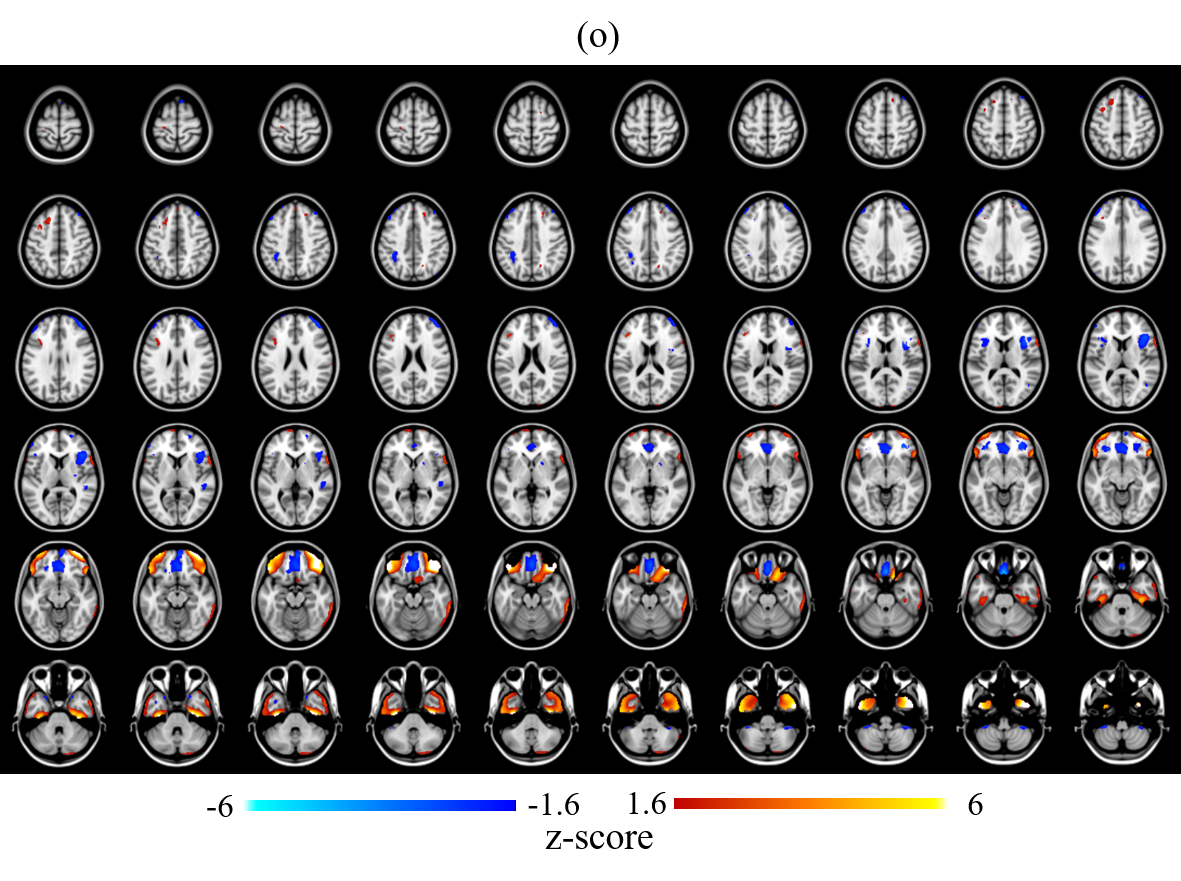

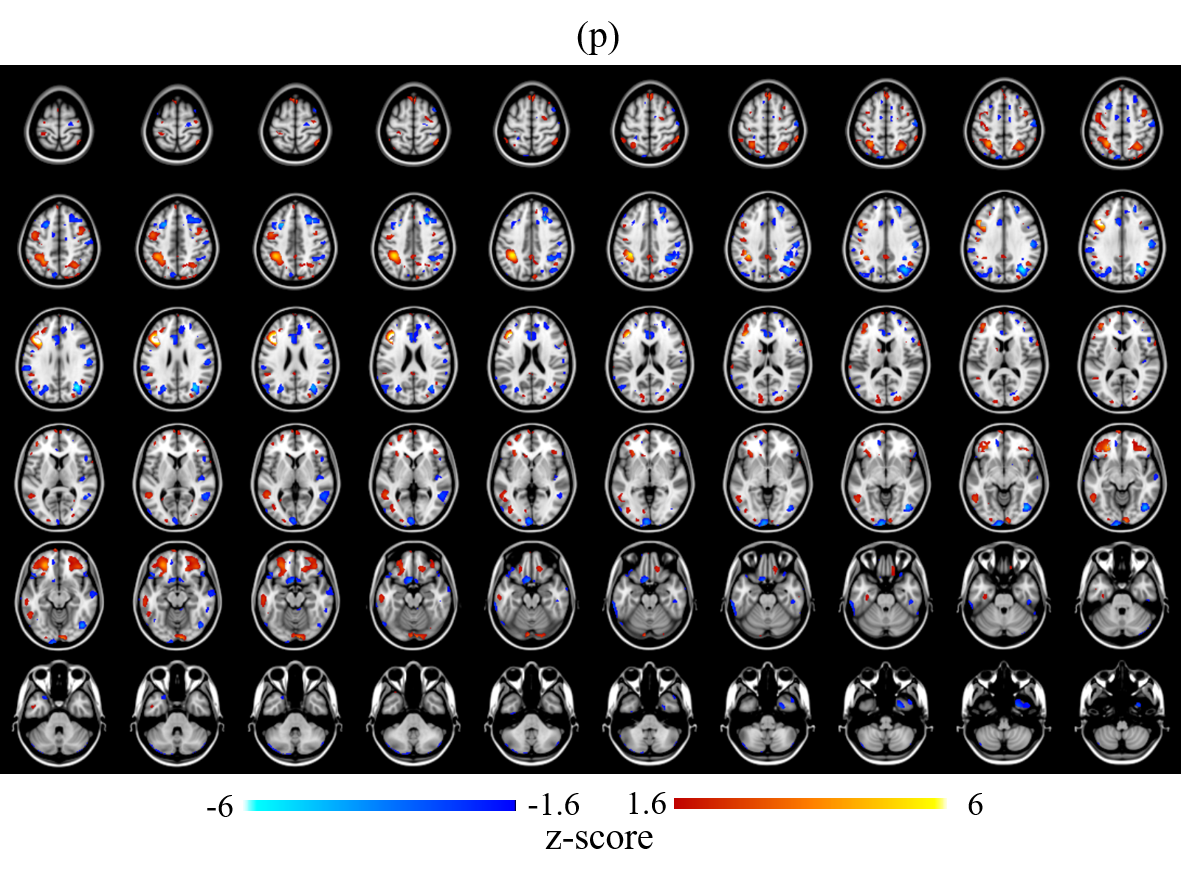

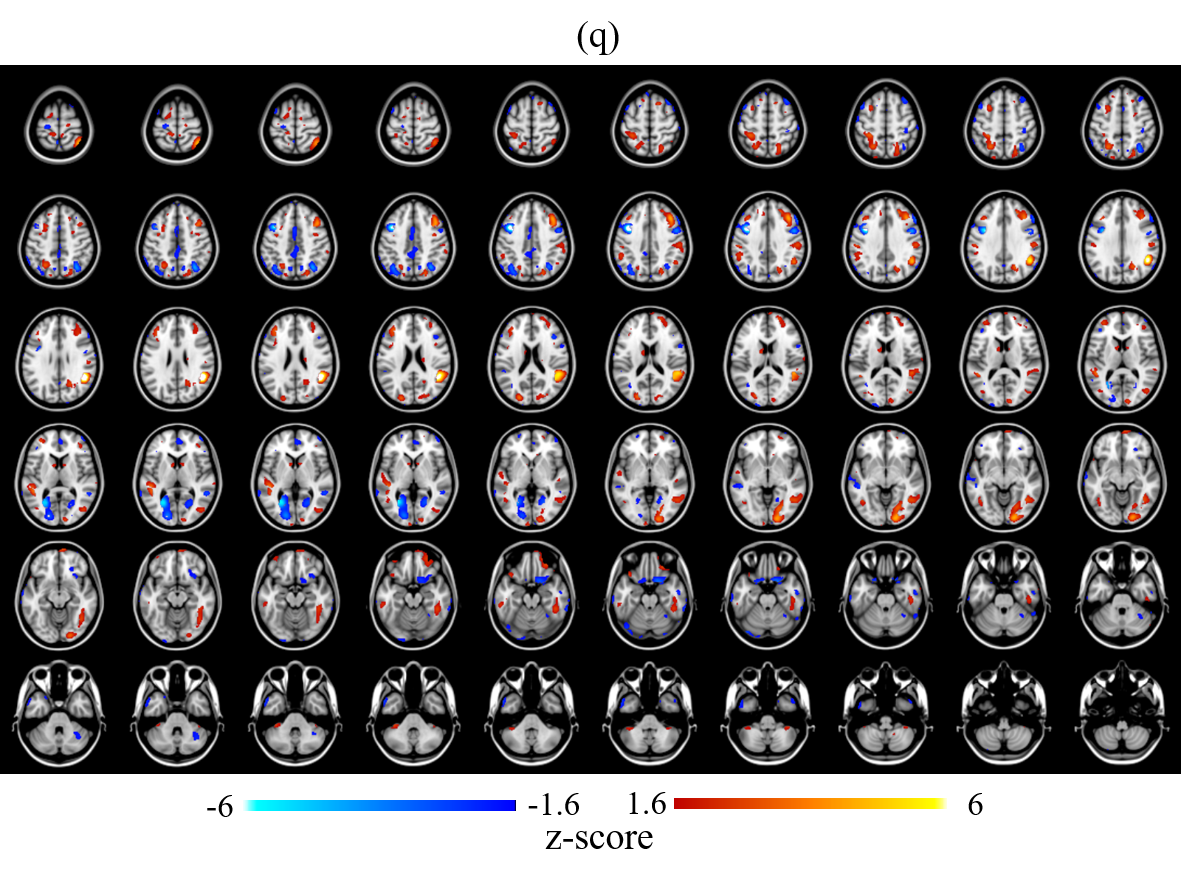

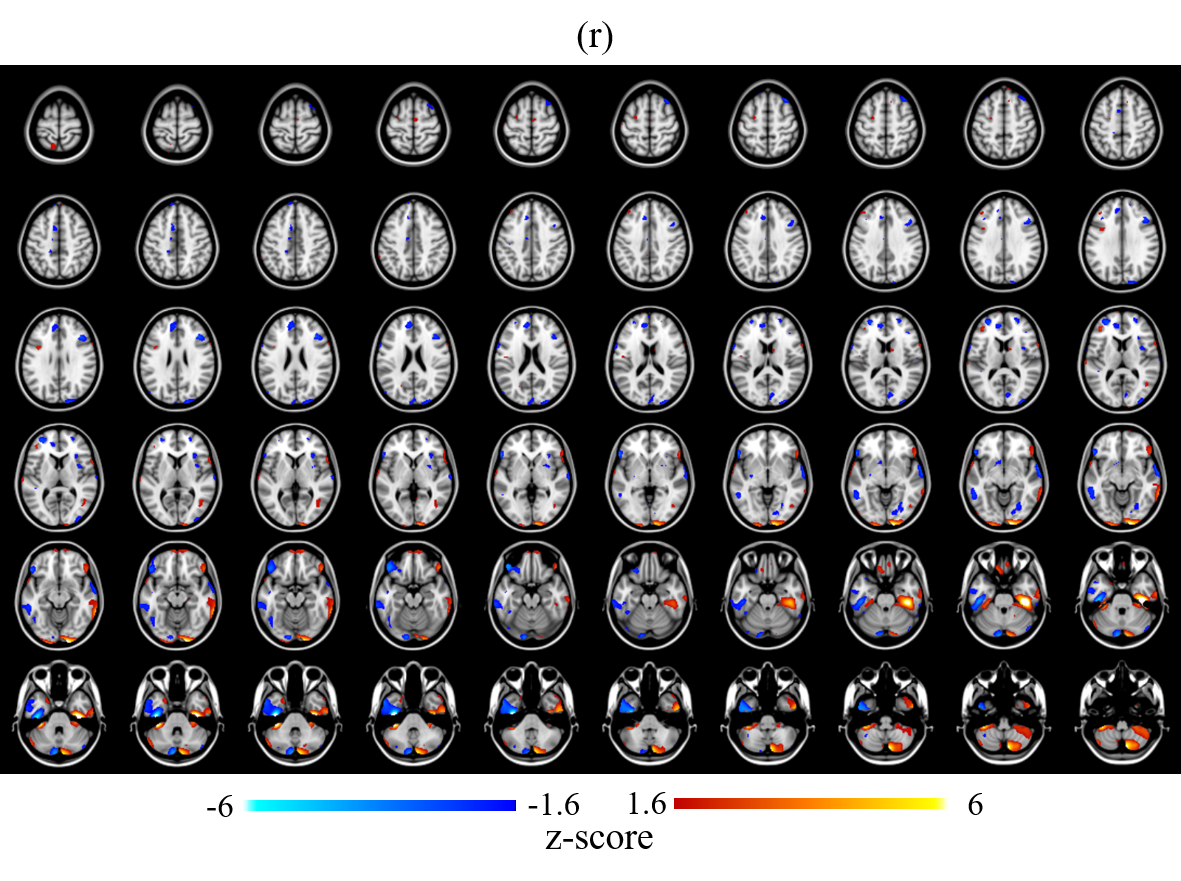

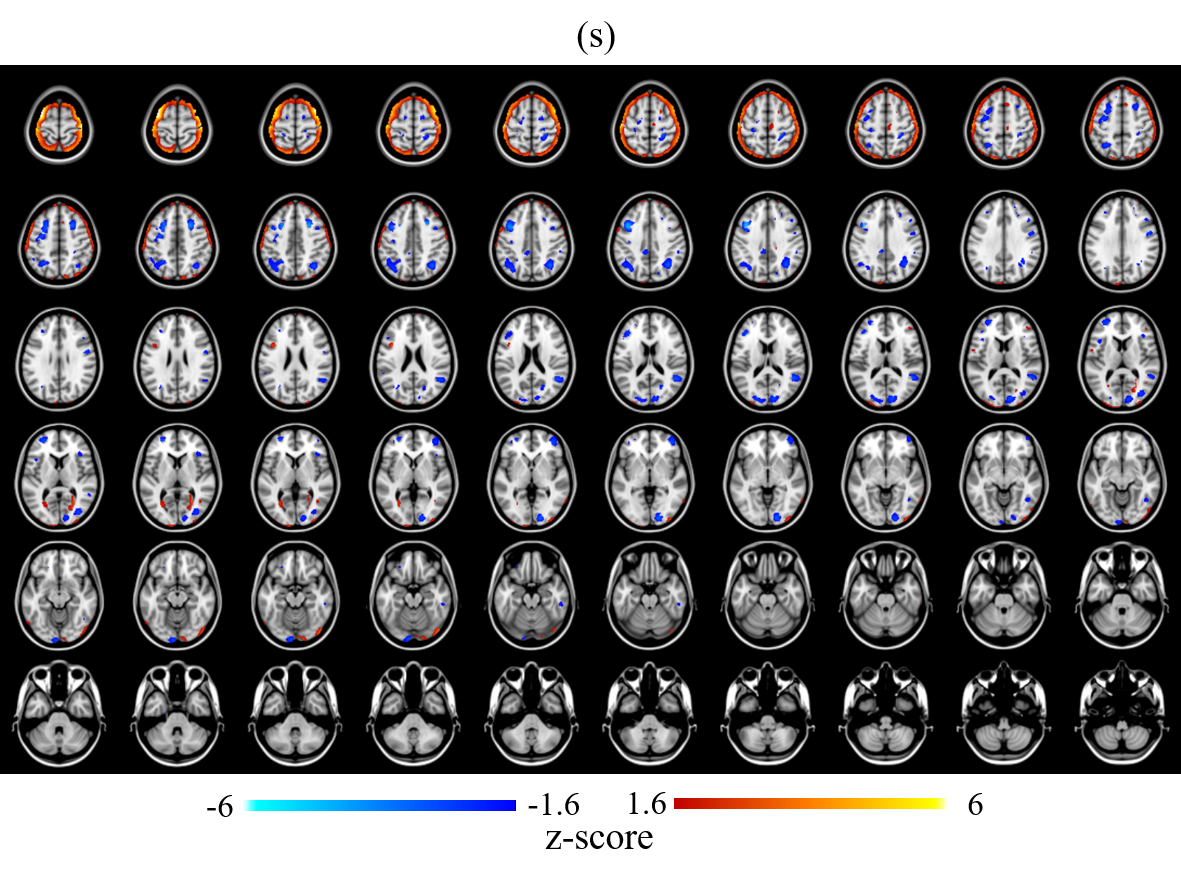

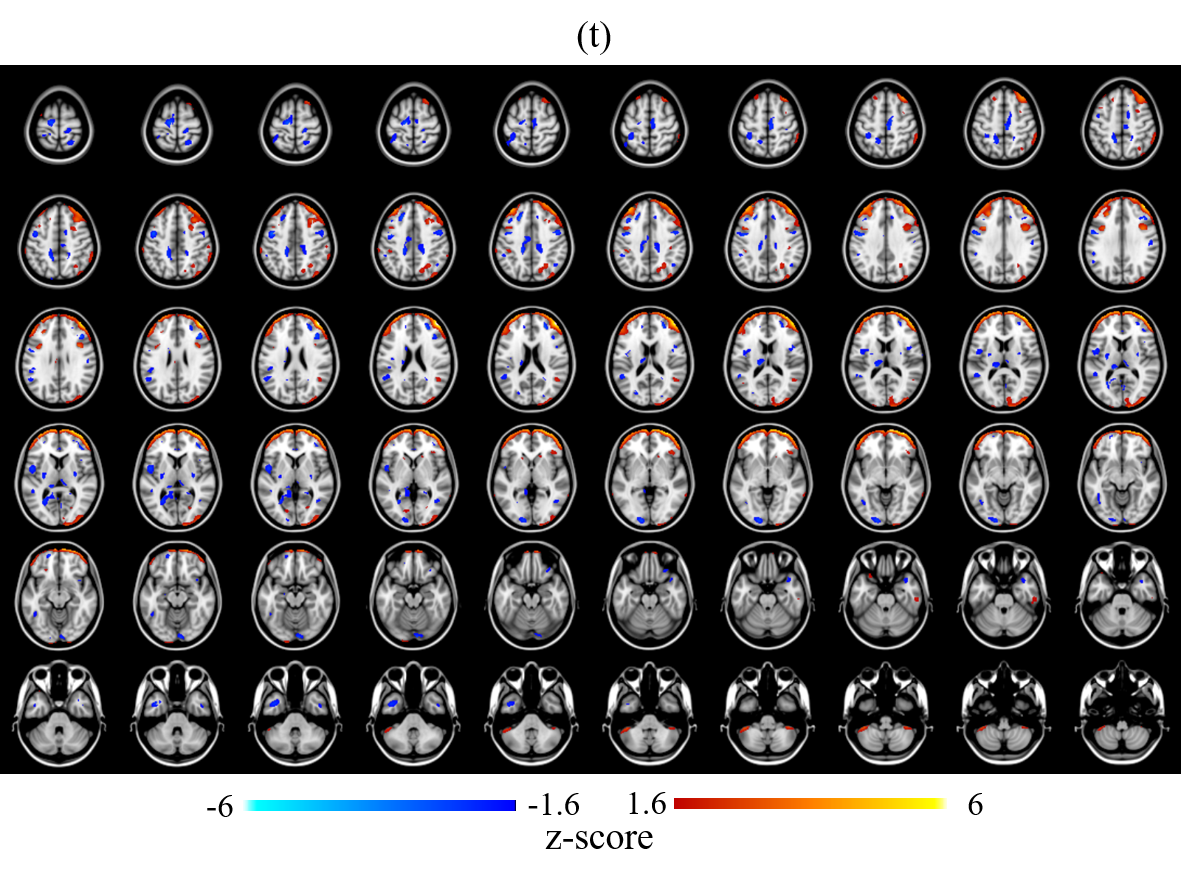
